# Population Genomics of Giant Mice from the Faroe Islands: Hybridization, Colonization, and a Novel Challenge to Identifying Genomic Targets of Selection

**DOI:** 10.1101/2025.01.20.633586

**Authors:** Bret A. Payseur, Peicheng Jing, Emma K. Howell, Megan E. Frayer, Eleanor P. Jones, Eyðfinn Magnussen, Jens-Kjeld Jensen, Yingguang Frank Chan, Jeremy B. Searle

**Affiliations:** Laboratory of Genetics, University of Wisconsin, Madison, WI 53706, USA; Fera Science, The National Agri-Food Innovation Campus, Sand Hutton, York YO41 1LZ, UK; School of Natural and Environmental Sciences, University of Newcastle, Newcastle NE1 7RU, UK; Faculty of Science and Technology, University of the Faroe Islands, Tórshavn, Faroe Islands; Naturalist, Í Geilini 37, FO-270 Nólsoy, Faroe Islands; Friedrich Miescher Laboratory of the Max Planck Society, 72076 Tübingen, Germany; Groningen Institute for Evolutionary Life Sciences (GELIFES), University of Groningen, 9747AG Groningen, The Netherlands; Department of Ecology and Evolutionary Biology, Cornell University, Ithaca, NY 14853, USA

**Keywords:** island evolution, hybridization, demographic inference, positive selection, house mouse

## Abstract

Populations that colonize islands provide unique insights into demography, adaptation, and the spread of invasive species. House mice on the Faroe Islands evolved exceptionally large bodies after colonization, generating interest from biologists since Darwin. To reconstruct the evolutionary history of these mice, we sequenced genomes of population samples from three Faroe Islands (Sandoy, Nólsoy, and Mykines) and Norway as a mainland comparison. Mice from the Faroe Islands are hybrids between the subspecies *Mus musculus domesticus* and *M. m. musculus*, with ancestry alternating along the genome. Analyses based on the site frequency spectrum of single nucleotide polymorphisms and the ancestral recombination graph (ARG) indicate that mice arrived on the Faroe Islands on a timescale consistent with transport by Norwegian Vikings, with colonization of Sandoy likely preceding colonization of Nólsoy. Substantial reductions in nucleotide diversity and effective population size associated with colonization suggest that mice on the Faroe Islands evolved large body size during periods of heightened genetic drift. Genomic scans for positive selection uncover windows with unusual site frequency spectra, but this pattern is mostly generated by clusters of singletons in individual mice. Variants showing evidence of selection in both Nólsoy and Sandoy based on the ARG are enriched for genes with neurological functions. Our findings reveal a dynamic evolutionary history for the enigmatic mice from Faroe Island and emphasize the challenges that accompany population genomic inferences in island populations.

**Significance Statement:** Populations that colonize islands are expected to have unusual histories compared to their mainland counterparts. Using population genomic data, we conclude that giant mice living on the Faroe Islands originated from hybrids, invaded the islands on a timescale consistent with transport by Vikings, and persisted despite drastic reductions in population size. We also uncover a novel challenge to scanning genomes for genes involved in adaptation.

## Introduction

The colonization of islands is a perilous task. To persist, populations that colonize islands must quickly adapt to novel environments in the face of drastic changes in population size and migration. As a result, island colonizers provide rich systems for studying the interplay of complex demography and natural selection (Losos and Ricklefs 2009). Evolutionary inferences from island populations also reveal conditions that favor the spread of invasive species, thereby informing conservation strategies (Estoup and Guillemaud 2010).

House mice (*Mus musculus*) are a model system for understanding island colonization on recent evolutionary timescales (Berry 1996). Among mammalian species, house mice are second only to humans in their global distribution (Angel et al. 2009). Human commensalism and a capacity for rapid adaptation has enabled house mice to invade islands around the world that encompass a wide variety of environments (Berry 1964; Berry and Peters 1975; Berry et al. 1978a; Berry et al. 1978b; Berry et al. 1979; Rowe-Rowe and Crafford 1992; Berry 1996; Berry 2009; Gabriel et al. 2010). Patterns of genetic variation in house mice can even be used to reconstruct human migration to islands (Jones et al. 2012; Jones et al. 2013).

One island radiation of house mice has captivated biologists since Darwin. The Faroe Islands comprise a small archipelago located in the North Atlantic approximately halfway between Norway and Iceland. The archipelago includes 18 islands, six of which are inhabited by house mice (Wilches et al. 2021). Mice on the Faroe Islands were cited as early examples of rapid adaptation and speciation by Darwin (1869), Degerbol (1942), and Huxley (1942). Based on the unusually large size of five specimens from Nólsoy, Clarke (1904) assigned mice from the Faroe Islands to a new subspecies (*Mus musculus faroensis*). Miller (1912) subsequently elevated distinction of the Faroe Island mice to the level of species. Later measurements confirmed the Faroe Island mice to be among the largest wild-living house mice in the world (Berry et al. 1978b). Due to cold temperatures on the Faroe Islands, mice there are suspected to be among the most climatically stressed in western Europe (Berry and Jakobson 1975), “probably approaching the physiological limit of their outdoor range” (Berry and Peters 1977). Mice from the Faroe Islands constitute one of several examples of house mice evolving extreme body size on islands (Berry 1964; Berry and Peters 1975; Berry et al. 1978a; Berry et al. 1978b; Rowe-Rowe and Crafford 1992). Genetic mapping revealed a complex genetic basis to size evolution, with 111 quantitative trait loci (QTL) that contribute to differences in body weight, body proportions, and growth hormone concentrations between mice from the Faroe Island of Mykines and a laboratory mouse strain selected for small body size (Wilches et al. 2021).

The first historical record of house mice inhabiting the Faroe Islands is in 1676 (Debes 1676). History (Edwards 2005), biogeography (Berry et al. 1978b), and mitochondrial sequence similarity (Jones et al. 2011) suggest that Norwegian Vikings (the Norse) were responsible for passively introducing mice to the Faroe Islands around 800 A.D., with potential later introductions of mice by the British, Dutch, German, and Danish (Jones et al. 2011). Based on analyses at a few molecular markers (Jones et al. 2011) and suspected origins in Norway, where many hybrids are found (Jones et al. 2010), it has been proposed that mice from the Faroe Islands derive ancestry from house mouse subspecies from western and eastern Europe (*M. musculus domesticus* and *M. m. musculus*, respectively) (Jones et al. 2011).

Key questions remain unanswered about the evolutionary history of the unusual mice living on the Faroe Islands. Are house mice on the Faroe Islands descended from hybrids? If so, how is the history of hybridization recorded in their genomes? When did house mice colonize the Faroe Islands? What population size changes accompanied island colonization? Did mice on the Faroe Islands continue to exchange genes with mice on the mainland after the island populations were established? Is there evidence for positive selection associated with island adaptation? The answers to these questions will reveal the conditions under which mice evolved extreme body sizes on the Faroe Islands.

Here, we take advantage of the unique perspective and expansive toolkit provided by population genomics to revisit the evolutionary history of the enigmatic house mice on the Faroe Islands, making comparisons with house mice from Norway as the colonization source. We reveal the genomic footprint of hybridization between subspecies and characterize the dynamics of demographic history associated with island colonization. We report genome-wide scans for positive selection that reveal a novel challenge with population genomic characterization of adaptation. Overall, we provide an increasingly rich evolutionary portrait of a fascinating, very recent island radiation.

## Results

### Genetic Differentiation between Populations

To visualize population structure among mice from the Faroe Islands, we conducted principal components analysis on the covariance matrix of genotypes (Price et al. 2006). Mice from Nólsoy, Sandoy, Mykines, and Norway form visually distinct groups based on their scores along principal components 1 and 2 (**Figure 1**). As a result, we treat mice from these geographic locations as separate populations in subsequent analyses. There also is evidence for population differentiation among mice within Norway, which we consider in the next section.

**Figure 1.**
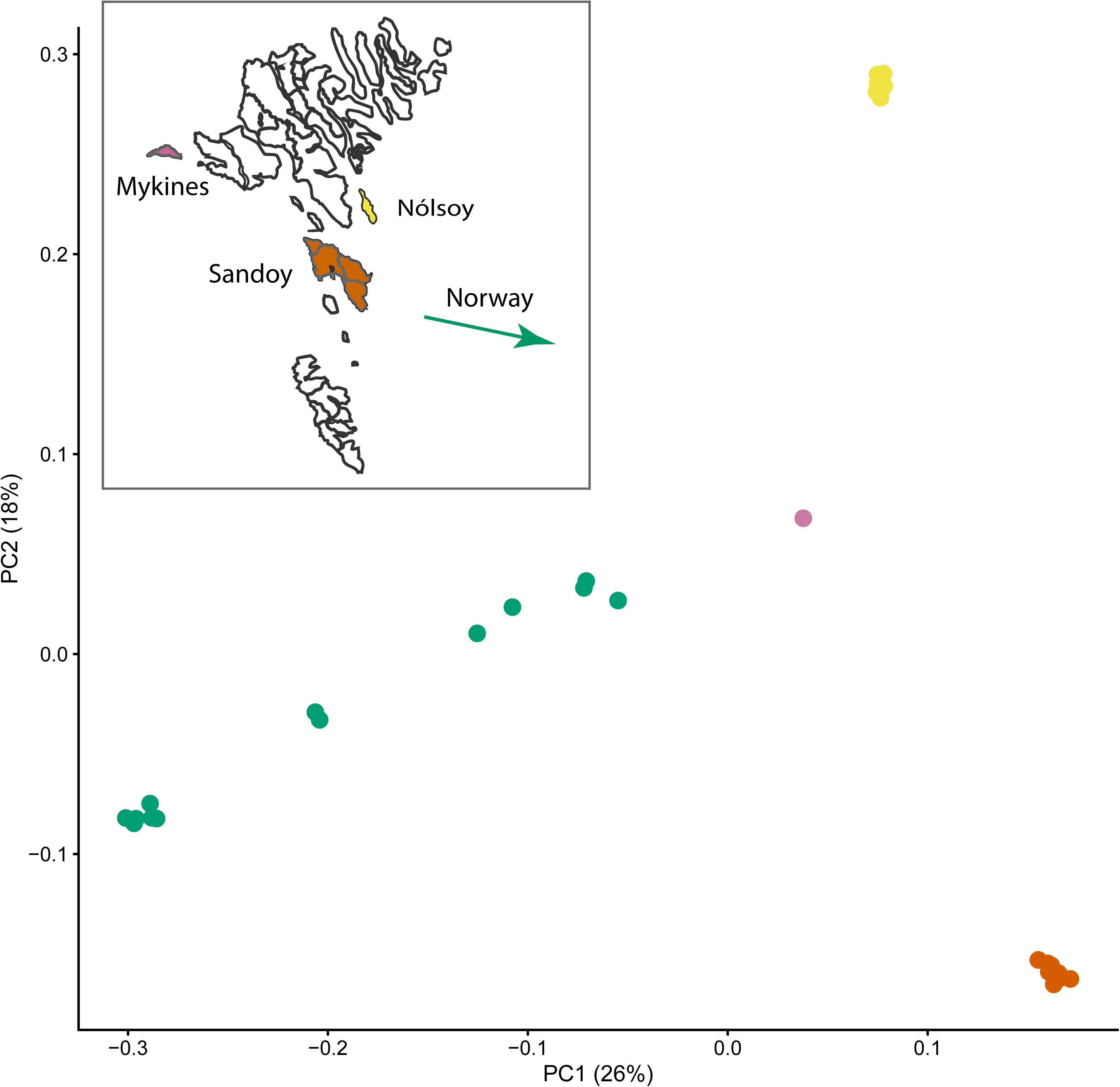
Genetic differentiation between mice from Nólsoy, Sandoy, Mykines, and Norway, and a map of the Faroe Islands. Each dot represents one mouse, plotted using its scores along principal components 1 and 2 inferred from principal components analysis of the genotype covariance matrix. Map of the Faroe Islands is inset, with an arrow marking the direction toward Norway. Color codes on the PCA plot match color codes on the map.

### Hybrid Ancestry

We used a probabilistic approach to infer the subspecific ancestries of Faroe Island mice (Corbett-Detig and Nielsen 2017). Ancestry analyses indicate that Faroe Island mice are hybrids between *M. m. domesticus* and *M. m. musculus* (*domesticus* and *musculus*, hereafter). Mice are inferred to be descended primarily from *domesticus* (**Figure 2**), with the genomic proportion of *domesticus* ancestry spanning 0.949 to 0.955 across Nólsoy mice, 0.906-0.920 across Sandoy mice, and 0.941 in one Mykines mouse (**Table S1**). Mice from Norway vary widely, with the genomic proportion of *domesticus* ancestry ranging from 0.095 to 0.925. Four mice from Norway with high proportions (> 0.9) of *domesticus* ancestry were used for analyses of island colonization history (see next section). Heterogenicity, the proportion of the genome heterozygous for ancestry, is 0.006-0.140 among mice from Nólsoy, 0.031-0.046 in mice from Sandoy, and 0.002 in one Mykines mouse. Heterogenicity values are higher in mice from Norway (0.024-0.259).

**Figure 2.**
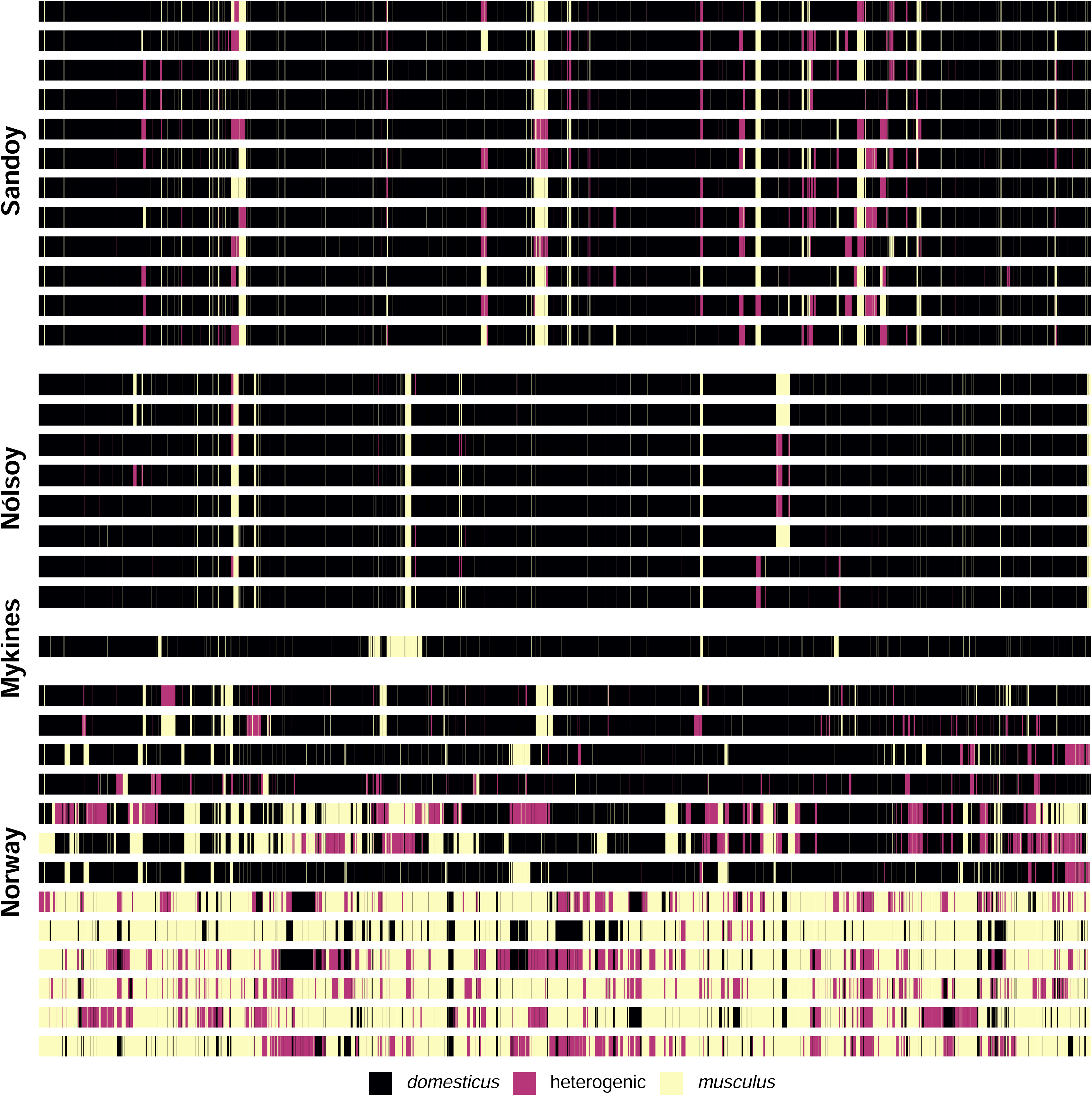
Subspecies ancestries along chromosome 3 for mice from Nólsoy, Sandoy, Mykines, and Norway. Each row is one mouse. Ancestries are color-coded. Ancestries for other chromosomes are shown in Figure S1.

Moving along a chromosome, subspecific ancestry alternates between *domesticus*, *musculus*, and heterogenic (ancestry from both subspecies) (**Figure 2**). Individual mice from Nólsoy, Sandoy, and Mykines harbor an average of 10.2, 10.3, and 10.6 junctions (switches) in ancestry per Mb, respectively. Junction numbers are higher in mice from Norway. There is variation in the spatial distribution of junctions within and among chromosomes (**Figure S1**). Junction patterns are more similar among island mice than among mainland mice from Norway.

Junctions represent historical recombination events that occurred in heterogenic regions of the genome (Baird 2006). To determine whether the rate of ancestry switching increases with recombination, we asked whether junction number is correlated with local recombination rate (cM/Mb; Cox et al. 2009) using comparisons across 10 Mb windows in mice from island populations. Spearman’s rank correlations between recombination rate and the average number of junctions across mice are *r* = 0.47 (*P* = 7.23 x 10^-13^) in Nólsoy, *r* = 0.47 (*P* = 6.78 x 10^-13^) in Sandoy, and *r* = 0.37 (*P* = 4.82 x 10^-8^) in Mykines.

The hybrid nature of the Faroe Island mice raises the possibility that body size evolution was partly shaped by subspecies ancestry. To test this hypothesis, we evaluated whether quantitative trait loci (QTL) for body size identified by crossing mice from Mykines with a laboratory strain (Wilches et al. 2021) are enriched for *domesticus* or *musculus* ancestry in 2 Mb windows in island populations. QTL intervals are enriched for *domesticus* ancestry in Nólsoy (Welch’s t-test; *t*_45_ = 4.55; *P* = 4.05 x 10^-5^), but not in Sandoy (*t*_25_ = −0.39; *P* = 0.7) nor in Mykines (*t*_25_ = −0.47; *P* = 0.64).

### Island Colonization History

#### LEVELS OF VARIATION

Focusing on all genomic regions with shared *domesticus* ancestry, mice on the Faroe Islands show substantial reductions in nucleotide diversity (the average number of pairwise differences; Tajima 1983). Average nucleotide diversity across windows is 0.066% in mice from Nólsoy, 0.078% in mice from Sandoy, and 0.199% in mice from Norway. Mice on islands harbor significantly less diversity than mice in Norway (paired *t*-tests; Nólsoy vs. Norway: *t*_322,778_ = −205.49, *P* < 2.2 x 10^-16^; Sandoy vs. Norway: *t*_289,431_ = −186.8, 2.2 x 10^-16^). The percentage of windows with zero diversity is 43.0% in mice from Nólsoy and 33.1% in mice from Sandoy, compared to 10.1% in mice from Norway. These low levels of genetic variation suggest that the history of Faroe Island mice involved reductions in population size.

#### THE SITE FREQUENCY SPECTRUM

To prepare for demographic analyses, we chose sequences with shared *domesticus* ancestry filtered to minimize selection at linked sites. Folded site frequency spectra (SFS) of single nucleotide polymorphisms (SNPs) residing within these regions differ among populations (**Figure 3**). Compared to SFS for mainland Norway, SFS for island populations (Nólsoy and Sandoy) display more evenness in the proportions of SNPs at common frequencies. Site frequency spectra for island populations also show bigger disparities between proportions of rare versus common SNPs. These characteristics of the island population SFS are consistent with non-equilibrium demographic histories, including changes in population size over time. Site frequency spectra also differ depending on whether they were estimated from genotype likelihoods (with ANGSD) or reconstructed from called genotypes (with GATK). For mice from Nólsoy, ANGSD estimates a much higher proportion of singleton SNPs than GATK. For mice from Sandoy, ANGSD estimates higher proportions of singletons and doubletons than GATK.

**Figure 3.**
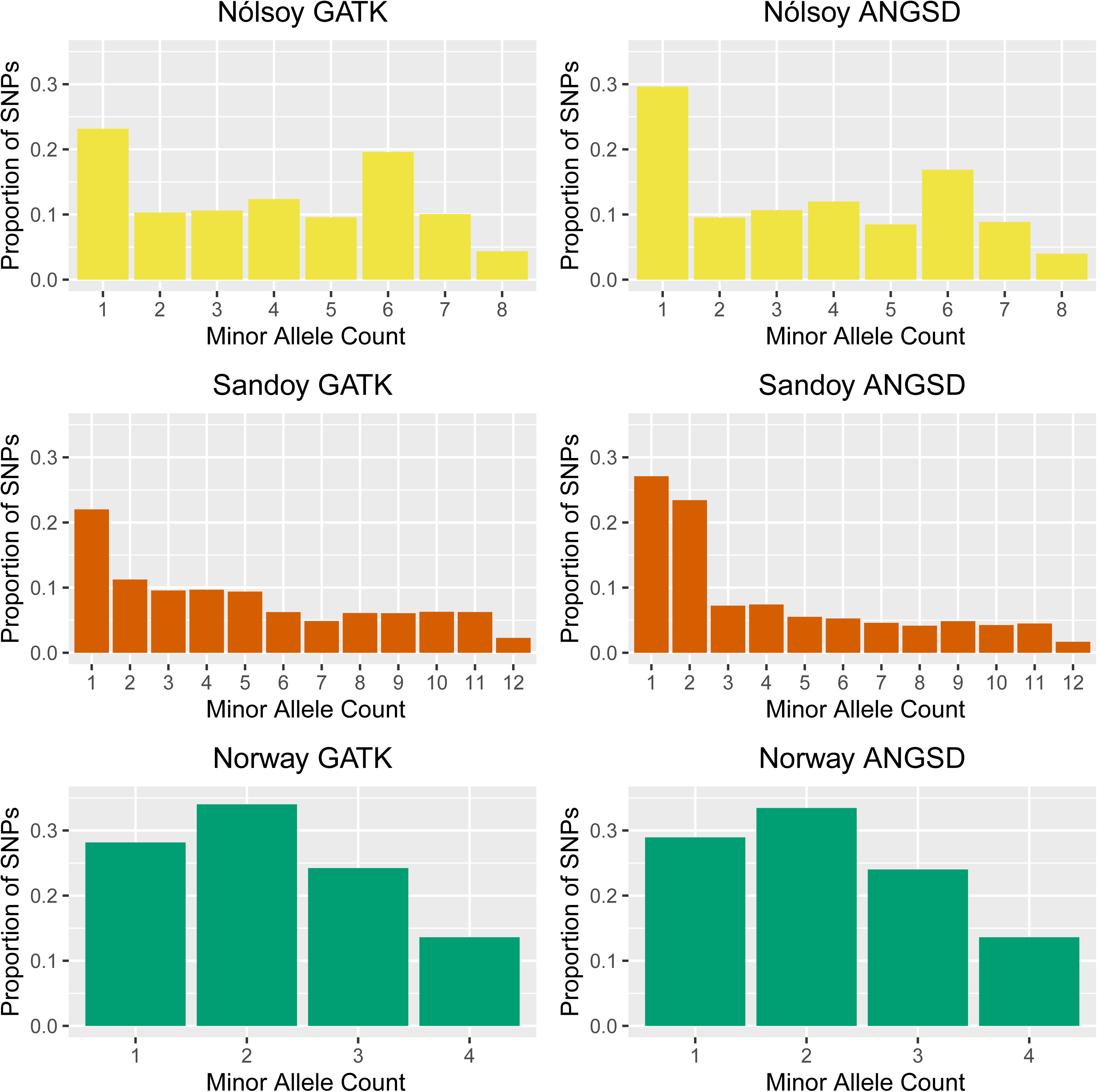
Site frequency spectra for Nólsoy, Sandoy, and Norway. Site frequency spectra were inferred from putatively neutral SNPs in genomic regions with shared *M. m. domesticus* ancestry, separately using GATK and ANGSD. Color-coding matches Figure 1 and Figure 4.

#### MODEL-BASED INFERENCES

To reconstruct demographic history associated with island colonization, we used two complementary approaches. In the first approach, we fit a variety of models to the folded, two-dimensional (2D) SFS (Gutenkunst et al. 2009) for three pairs of populations: Norway-Sandoy, Norway-Nólsoy, and Sandoy-Nólsoy. In light of differences between the SFS recovered with GATK and ANGSD, we conducted separate sets of demographic analyses on the 2D SFS inferred by each method.

In the second approach, we estimated demographic parameters from ancestral recombination graphs for each pair of populations (ARG) (Speidel et al. 2019). In contrast to SFS-based analyses, ARG-based analyses incorporated variation throughout the genome, regardless of subspecies ancestry, used only GATK-derived genotypes, and focused on derived alleles inferred using two outgroup species (*M. caroli* and *M. spretus*). Here, we highlight comparisons between estimates of the same demographic parameters from the two approaches.

#### NORWAY-SANDOY

The best-fitting demographic model for the Norway-Sandoy GATK-derived 2D SFS includes a population size reduction in Norway followed by a severe bottleneck in Sandoy after its split from Norway (**Figure 4**). The best-fitting model for the Norway-Sandoy ANGSD-derived 2D SFS includes these features plus one-way migration from Norway to Sandoy (**Figure S2**). For some parameter estimates, disparities are large between GATK-derived SFS and ANGSD-derived SFS (**Table S2**). Effective population size (*N_e_*) estimates for Sandoy immediately following the split from mainland Norway are 760 (GATK) and 7 (ANGSD). The timing of the population split ranges from 3,176 (GATK) to 5,411 (ANGSD) generations ago. The Sandoy bottleneck is estimated to end 1,286 (GATK) or 5,396 (ANGSD) generations ago. Both best-fitting models include historical inbreeding (**Table S2**).

**Figure 4.**
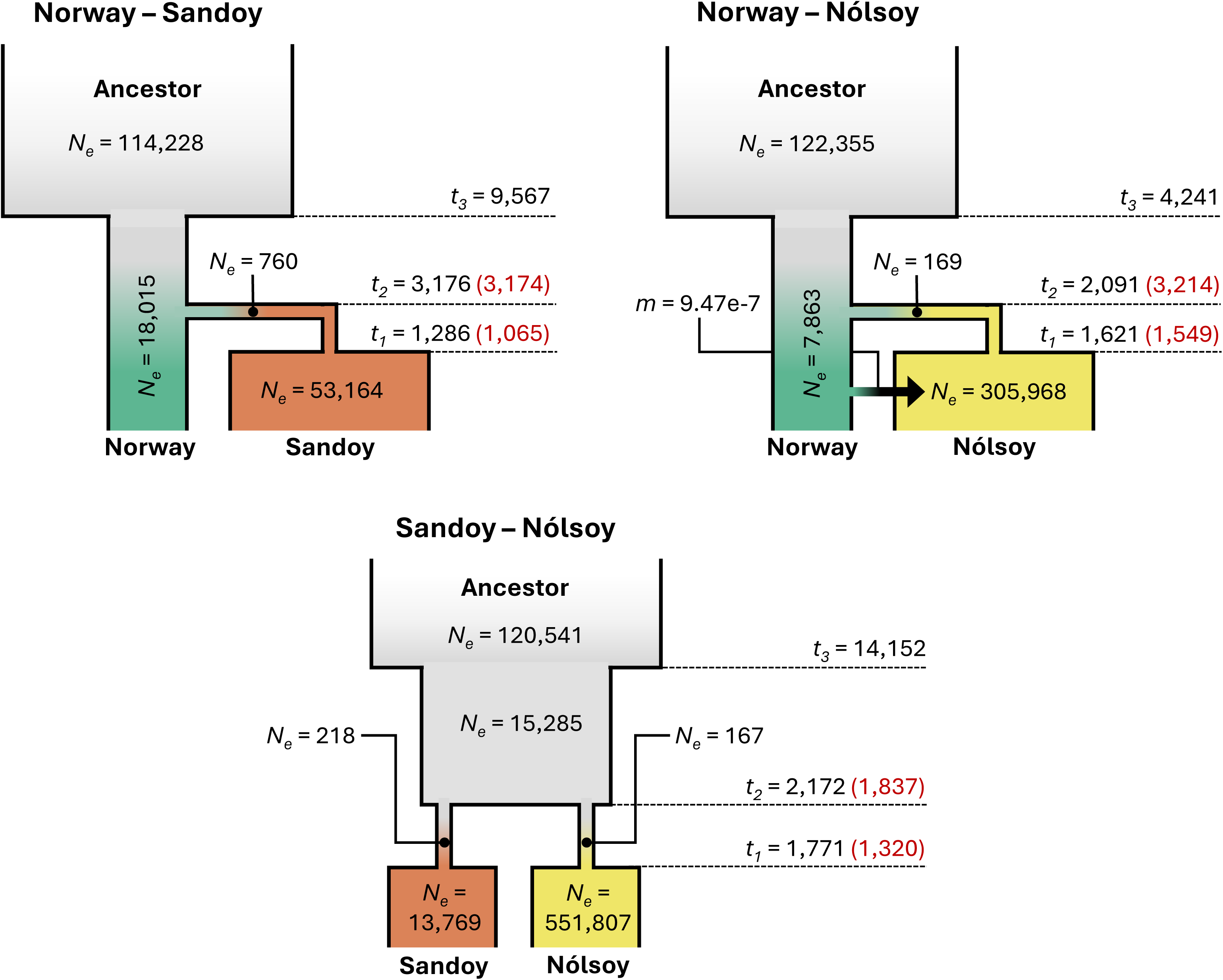
Best-fitting demographic models for mice from Nólsoy and Sandoy, treating mice from Norway as a mainland reference population. Parameter estimates come from analyses of two-dimensional site frequency spectra for pairs of populations inferred from GATK-derived genotypes for putatively neutral SNPs in genomic regions with shared *M. m. domesticus* ancestry. Estimates for comparable parameters from analyses of ancestral recombination graphs are denoted in red. *N_e_* = effective population size. *t* = timing in generations of population splits or changes in *N_e_*, ordered beginning with the most recent event. *m* = migration rate.

The split time estimate for Norway-Sandoy from the ARG is 3,174 generations ago, very close to the corresponding parameter estimate from the GATK-derived SFS of 3,176 generations ago (**Figure 4**). For Sandoy, the ARG yields a minimum *N_e_* at 1,065 generations ago (**Figure S3**), which is close to the end of the bottleneck from the GATK-derived SFS at 1,286 generations ago. The ARG estimate for lowest Sandoy *N_e_* is 1,178, compared to the GATK-derived SFS *N_e_* of 760 immediately after the split from mainland Norway.

#### NORWAY-NÓLSOY

Similar to the Norway-Sandoy comparison, the best-fitting demographic model for the Norway-Nólsoy GATK-derived 2D SFS includes a population size reduction in Norway followed by a bottleneck in Nólsoy that accompanies the population split from the mainland. The best-fitting GATK-derived model additionally includes one-way migration from Norway to Nólsoy after their split (**Figure 4**). The pre-split mainland size reduction is absent from the best-fitting model for the Norway-Nólsoy ANGSD-derived 2D SFS (**Figure S2**). Disparities in parameter estimates between GATK-derived SFS and ANGSD-derived SFS are smaller for Norway-Nólsoy than for Norway-Sandoy. For both models, *N_e_* of Nólsoy is small (169 vs. 9) immediately following its split from mainland Norway and *N_e_* recovers soon afterward, suggesting rapid expansion of island mice. The split time between Norway and Nólsoy is estimated to be 2,091 (GATK-derived SFS) or 3,532 (ANGSD-derived SFS) generations ago. Both models support historical inbreeding and a low rate of migration from Norway to Nólsoy (**Table S2**). Modest standard deviations accompany most parameter estimates (**Table S2**).

The Nólsoy-Norway split time is estimated from the ARG to be 3,214 generations ago, which can be compared to the time of 2,091 generations ago estimated from the GATK-derived SFS (**Figure 4**). For Nólsoy, the minimum *N_e_* reconstructed from the ARG occurs 1,549 generations ago, which is close to the time of 1,621 generations ago at which the bottleneck ends, as estimated from the GATK-derived SFS. The ARG estimate for the lowest Nólsoy *N_e_* is 1,183 (**Figure S3**), compared to 169 from the GATK-derived SFS.

#### SANDOY-NÓLSOY

The best-fitting model for the Sandoy-Nólsoy GATK-derived 2D SFS includes a reduction in ancestral population size followed by a bottleneck on both islands (**Figure 4**). The best-fitting model for the Sandoy-Nólsoy ANGSD-derived 2D SFS excludes the ancestral size reduction and includes migration from Nólsoy to Sandoy following their split (**Figure S2**). *N_e_* estimates are similar for Sandoy and Nólsoy for both GATK-derived SFS and ANGSD-derived SFS (**Table S2**). Population split time is 2,172 (GATK) or 3,774 (ANGSD) generations ago. Both models include historical inbreeding (**Table S2**).

The split time for the two island populations based on the ARG (1,837 generations ago) is close to the split time inferred from the GATK-derived SFS (2,172 generations ago) (**Figure 4**). The times with the lowest *N_e_* are similar for Sandoy (1,320 generations ago) and Nólsoy (1,349 generations ago) (**Figure S3**), and a few hundred generations more recent than the bottleneck end estimated from the GATK-derived SFS (1,771). The ARG estimate for lowest island *N_e_* is 955, compared to *N_e_* following the population split based on the GATK-derived SFS (Sandoy: 218; Nólsoy: 167).

### Genomic Scans for Selection Connected to Island Adaptation

#### SELECTION SCANS BASED ON SUMMARY STATISTICS

To generate expected patterns of variation under neutrality, we conducted coalescent simulations assuming parameter estimates from models without inbreeding that showed the best fit to the GATK-derived SFS. Distributions of nucleotide diversity, Tajima’s D (a measure of skew in the SFS; Tajima 1989), and F_ST_ (a measure of population differentiation; Hudson et al. 1992) computed from windows in datasets simulated under these models are close to distributions from the real datasets including all windows with shared *domesticus* ancestry (**Figure S4**), suggesting they provide reasonable null models for genomic scans for selection.

Our first genomic scan for selection considered metrics that summarize the SFS (Tajima’s D) and population differentiation (F_ST_) between island and mainland. Across windows with shared *domesticus* ancestry, 84 windows in Nólsoy and 57 windows in Sandoy have Tajima’s D values that are unexpectedly negative for neutrality (i.e. in the direction expected under a simple selective sweep; Braverman et al. 1995) (**Table S3**), given simulated demographic histories and strict corrections for multiple testing (genome-wide Bonferroni *P* < 0.05). Tajima’s D values in these windows range from −2.52 to −2.43 in Nólsoy and range from −2.75 to −2.62 in Sandoy.

As expected, large percentages of the variants in these windows with unusual Tajima’s D values are singletons (96.6-100% in Nólsoy; 96.1-100% in Sandoy). Surprisingly, all or most of the singletons in these windows are found in a single mouse, with the identity of the mouse varying among windows (**Table S3**). This observation comes from a non-trivial number of singletons in these outlier windows (Nólsoy average: 59.2 singletons; Sandoy average: 64.5 singletons), so it is not a simple consequence of low levels of variation. This pattern seems incompatible with positive selection as the cause of the unusual values of Tajima’s D in these windows, so we do not pursue them further. No windows show unusual values of F_ST_ once demographic history and multiple testing are accounted for.

#### SELECTION SCANS BASED ON THE FULL SITE FREQUENCY SPECTRUM

Our second genomic scan for selection searched for windows with SFS that depart from the genome-wide SFS (De Georgio et al. 2016) in shared *domesticus* regions. Two thousand seven hundred and forty-six windows in Nólsoy and 154 windows in Sandoy have likelihood ratio test statistics that are unusual for neutrality, given the demographic history and strict corrections for multiple testing (genome-wide Bonferroni *P* < 0.05) (**Table S4**). None of these windows are the same in Nólsoy and Sandoy. These windows with unusual SFS form several clusters in the genome.

Although this approach detects more windows with potential evidence of positive selection than scans based on Tajima’s D (above), most of these windows are afflicted by the same issue: an excess of singletons derived mostly or exclusively from a single mouse, with the identity of this mouse varying among windows (**Table S4**). Since this pattern seems unexpected under models of positive selection, we do not pursue these windows further.

#### SELECTION SCANS BASED ON THE ANCESTRAL RECOMBINATION GRAPH

Our third approach to scanning the genome for selection searched for derived SNPs with unusually high frequencies given their genealogies, as inferred using the ARG (Speidel et al. 2019). In contrast to scans based on the SFS, this approach incorporated regions throughout the genome, irrespective of subspecies ancestry. Although ARG analyses identify SNPs with evidence for positive selection, none of the tests survive corrections for multiple testing based on either the number of SNPs or the estimated number of genealogies across the genome. A total of 670 SNPs have uncorrected *P* < 0.05 in both Nólsoy and Sandoy, but not in Norway (**Table S5**). These SNPs are clustered into a modest number of genomic regions. Genes in these intervals are significantly enriched for biological processes and cellular components connected to neurological features and functions, including synapses between neurons and synaptic vesicle fusion (**Table S5**).

## Discussion

Mice living on the Faroe Islands have complex evolutionary histories. As suspected based on previous findings (Jones et al. 2011), they are products of hybridization between *domesticus* and *musculus*. Faroe Island mice are primarily *domesticus* in origin, probably reflecting a history of repeated backcrossing to *domesticus* following initial hybridization. The skew toward *domesticus* ancestry matches the morphological resemblance of Faroe Island mice to this subspecies (Jones et al. 2011). The hybrid nature of Faroe Island mice is likely connected to ancestors in Norway, where our genomic ancestry inferences support earlier claims of a hybridization gradient (Jones et al. 2010). Although firm conclusions await the application of methods that consider a range of hybridization histories, the overall similarity in junction patterns among island mice, as well as between island mice and some Norway mice, suggests that the hybridization visible in Faroe Island mouse genomes occurred prior to island colonization. We uncovered signs that subspecific ancestry could be connected to the evolution of body size. QTL contributing to body weight differences between Faroe Island mice and laboratory mice (Wilches et al. 2021) are enriched for *domesticus* ancestry, at least in mice from Nólsoy.

To reconstruct colonization histories, we identified best-fit models based on separate analyses of the ANGSD-derived 2D SFS in genomic regions with *domesticus* ancestry, the GATK-derived 2D SFS in genomic regions with *domesticus* ancestry, and the ancestral recombination graph across the genome. Considering ranges of point estimates across the three approaches and assuming three generations per year (Berry et al. 2008) indicates that mice colonized Nólsoy 697-1,071 years ago and Sandoy 1,058-1,804 years ago. The split time between Nólsoy and Sandoy is estimated to be 724-1,258 years ago. Collectively, these results suggest that mice colonized Sandoy first and Nólsoy later. Overall, the timing of colonization is consistent with passive transport of mice to the Faroe Islands by Norwegian Vikings (Edwards 2005; Jones et al. 2011). Mice on these islands likely had several hundred to a few thousand generations following colonization to evolve unusually large bodies.

Mice on the Faroe Islands appear to have persisted through substantial declines in effective population size (*N_e_*). Estimates of *N_e_* associated with island colonization are as low as 2 and the island populations show low levels of nucleotide diversity. In populations with temporal variation in number, *N_e_* is expected to be influenced disproportionately by intervals with small numbers. Consequently, low *N_e_* values associated with colonization raise the prospect of heightened genetic drift and reduced selective efficacy during a substantial proportion of the time mice have occupied the Faroe Islands. In turn, these inferences should motivate careful evaluation of the relative roles of natural selection and genetic drift in the evolution of extreme body size and other phenotypes in these mice.

We uncovered evidence for limited migration following island colonization. Best-fit models to the 2D SFS feature low rates of migration from Norway to Nólsoy (both ANGSD-derived SFS and GATK-derived SFS) and from Norway to Sandoy (ANGSD-derived SFS). Restrictions to gene flow may have facilitated the evolution of genetic and phenotypic differences between these groups (Berry et al. 1978; Davis 1983; Jones et al. 2011). In addition, our results suggest that mice on the Faroe Islands and in Norway have a history of inbreeding. This inference matches the conclusion from a wider survey of house mice based on microarray genotypes, which uncovered widespread and variable evidence of inbreeding across populations (Morgan et al. 2022).

Our study underscores challenges that accompany evolutionary inferences from genome sequences in populations with complex demographic histories. The characterization of demographic history can depend on which methods are used to reconstruct the SFS. In our study, discordance between best-fit models from ANGSD-derived SFS vs. GATK-derived SFS persisted despite decent sequencing coverage and the application of best practices for these popular approaches. Although estimates of key parameters are broadly similar between the two methods, our results provide a reminder that uncertainty in the data should be incorporated during population genomic analyses (Johri et al. 2022). Consideration of the model-based population genetic methods we used to reconstruct colonization histories also emphasizes the role of uncertainty. Some parameter estimates of best-fit models from analyses of the SFS feature non-trivial confidence intervals, and the ARG method implemented in *Relate* currently yields only point estimates. Furthermore, samples of wild populations can contain high incidences of close relatives. The modest sample sizes that remained after we excluded close relatives represent additional sources of error. Based on our experience, we recommend that population geneticists examine relatedness before proceeding to reconstruct demographic history.

Another lesson of general interest from our investigation concerns the goal of identifying genomic windows that contain beneficial mutations involved in adaptation. Scanning the local SFS in *domesticus* regions pinpointed many windows showing conventional evidence of positive selection even after accounting for demographic history and multiple testing. Interestingly, most of these windows harbor an excess of singletons contributed by a single mouse (with the identity of the mouse varying among windows). This pattern contrasts with expectations for a selective sweep, in which singletons should accumulate randomly across individuals following the loss of neutral variation caused by fixation of a linked beneficial variant. We confirmed that mice harboring clusters of singletons in these windows have high-confidence ancestry calls, arguing against the explanation that most of the clusters are caused by mis-inference of heterogenic mice as carrying homozygous *domesticus* ancestry. Whatever explains this constellation of singletons, it could be a source of false positives in selection scans that is currently unrecognized in the field and deserves further examination. We recommend that researchers using the SFS to search for positive selection inspect the partitioning of variants among individuals. Overall, our results underscore the barriers to localizing targets of adaptive evolution in the genomes of recently bottlenecked populations (Barton 2000; Poh et al. 2014), including populations that colonize islands.

The ARG offers strategies for characterizing positive selection that complement those based on traditional population genetic approaches. SNPs with low *p*-values in both Nólsoy and Sandoy given their local genealogies may reflect parallel adaptation of mice on these two islands. The enrichment of genes with neurological functions we detected raises the prospect that mice evolved new behaviors following their colonization of the Faroe Islands. Mice from Gough Island evolved increased exploration and boldness (Stratton et al. 2021; Stratton et al. 2023), and genes in genomic regions showing evidence of positive selection in these mice are enriched for neurological functions (Payseur and Jing 2021). Still, gene ontology results from Faroe Island mice should be viewed with caution since they come from applying relaxed significance thresholds. In addition, the effects of errors in genealogy reconstruction on ARG-based tests for selection deserve further examination.

## Materials and Methods

### Mice

Mice were collected on Sandoy (from 5 localities) during 2005-2006, and on the small islands of Mykines (October 2009) and Nólsoy (between 1984 and 2003). Because of the small sizes of the villages on Nólsoy and Mykines, sampling locations were less than 1km apart and treated as a single locality. DNA from island mice was extracted from tail or ear clips. Tissues were lysed with lysis buffer with Tris-EDTA-SDS, SDS and protein were precipitated using potassium acetate, and DNA was purified using magnetic beads. In Norway, mice were collected from five geographical districts from 8/31/2007 to 10/13/2007: Sogn og Fjordane, Hordaland, Sogn, Vestfold and Telemark. DNA from Norway mice was extracted from muscle using the Qiagen Puregene kit.

### Genome Sequencing, Alignment, Variant Calling, and Filtering

We sequenced the genomes of 15 mice from Nólsoy, 20 mice from Sandoy, 7 mice from Mykines, and 15 mice from Norway (7 from Telemark, 2 from Sogn og Fjordane, 2 from Hordaland, 2 from Sogn, and 2 from Vestfold). Sequencing was completed by the Biotechnology Center at the University of Wisconsin-Madison. Genomic DNA concentrations were verified using the Qubit® dsDNA HS Assay Kit (Life Technologies, Grand Island, NY). Samples were prepared according to the Celero PCR Workflow with Enzymatic Fragmentation (Tecan Genomics, Redwood City, CA). Quality and quantity of the finished libraries were assessed using an Agilent Tapestation (Agilent, Santa Clara, CA) and Qubit® dsDNA HS Assay Kit, respectively. Paired end, 150 bp sequences were generated using the Illumina NovaSeq6000 (Illumina, San Diego, CA). Sequencing reads from genome sequences for 8 mice from Germany, 8 mice from France, 8 mice from Czech Republic, and 8 mice from Kazakhstan (Harr et al. 2016) were downloaded from: ftp.sra.ebi.ac.uk/vol1/ERA441/ERA441366/fastq/; ftp.sra.ebi.ac.uk/vol1/ERA532/ERA532416/fastq/; ftp://ftp.sra.ebi.ac.uk/vol1/run/ERR112.

Sequencing reads were aligned to the mouse mm10 genome reference sequence downloaded from the UCSC Genome Browser (Kent et al. 2002), using bwa-mem (Li and Durbin 2009) with default settings. Bam files were sorted and merged using Samtools (Danecek et al. 2021). Duplicates were marked and removed with Picard tools (https://broadinstitute.github.io/picard/). Indel realignment was performed using IndelRealigner in GATK. Variants were identified and genotyped using GATK (McKenna et al. 2010). Average realized sequencing coverage was close to the target of 20X (**Table S6**). gvcf files generated by HaplotypeCaller (GATK) were merged into a joint .vcf file using GenotypeGVCFs (GATK), and SNPs were processed with VariantFiltration following filtering criteria provided by GATK guidelines for best practices. Genotypes were called separately for each population.

Close relatives were identified by pairwise identity-by-state comparisons with KING (Manichaikul et al. 2010) using the ‘kinship’ option on all SNPs that passed GATK filters (results using SNPs pruned for linkage disequilibrium were similar). Prior to downstream analyses, mice inferred to be close relatives were removed. For each population, we chose unrelated mice by starting with the set selected using the KING ‘unrelated’ option (which removed first-degree and second-degree relatives), adjusting it to leave no third-degree relatives and to preferentially remove the mouse with lower sequencing coverage from each relative pair. This approach left 8 mice from Nólsoy, 12 mice from Sandoy, and 13 mice from Norway (7 from Telemark, 2 from Hordaland, 2 from Sogn, 1 from Sogn og Fjordane, and 1 from Vestfold). Following the same procedure left unrelated samples of 8 mice from Germany, 4 from France, 7 from Czech Republic and 4 from Kazakhstan. All 7 mice from Mykines were inferred to be first-degree relatives. As a result, we did not attempt to reconstruct colonization history for Mykines, though we inferred the subspecific ancestry of a single mouse from Mykines. Subsequent analyses only considered these samples of unrelated mice; genotypes were recalled after closely related mice were removed.

Genotypes were merged across populations with Bcftools (Danacek et al. 2021). Positions with missing genotypes were identified in each population and filled in using the ‘all-sites’ option in GATK. SNPs within repetitive sequences and segmental duplications annotated for the reference genome were removed with Bedtools (Quinlan et al. 2010) ‘subtract’ using positions from ‘RepeatMasker’ and ‘Segmental Dups’ tracks downloaded from the UCSC Genome Browser.

All analyses focused on SNPs located on the autosomes.

### Inference of Genetic Differentiation and Ancestry

Genetic differentiation between mice from different locations was visually examined using principal component analysis (PCA) of the genotype covariance matrix, as implemented in SmartPCA (Price et al. 2006). Prior to PCA, SNPs with minor allele frequency less than 0.05, greater than 50% missing genotypes, and/or pairwise *R^2^* higher than 0.2 were removed.

We used AncestryHMM (Corbett-Detig and Nielsen 2017) to probabilistically assign subspecific ancestry (homozygous *domesticus*, homozygous *musculus*, and heterozygous *domesticus/musculus*) along the genome of individual mice from Nólsoy, Sandoy, Mykines, and Norway. We computed recombination distances between pairs of adjacent SNPs by interpolation from their physical distances and local recombination rates taken from the mouse genetic map constructed from the heterogeneous stock (Cox et al. 2009; downloaded at https://github.com/kbroman/CoxMapV3). We treated mice from France and Germany as *domesticus* reference populations and treated mice from Czech Republic and Kazakhstan as *musculus* reference populations. We ran separate sets of AncestryHMM analyses, treating different intersubspecific pairs of populations as references (France-Czech, France-Kazakhstan, Germany-Czech, and Germany-Kazakhstan). To improve the quality of inferred ancestry calls, we used two conservative approaches. First, we only considered SNPs for which one of the three possible ancestries was assigned a posterior probability > 0.95. Second, we required that ancestry calls agreed across analyses using each of the four pairs of reference populations.

We created plots to visualize ancestry along each chromosome. We also used a few metrics to summarize inferred patterns of ancestry. For each mouse, we computed the proportion of each chromosome with *domesticus* ancestry. Heterogenicity was calculated as the proportion of each chromosome in each mouse inferred to harbor ancestry from both *domesticus* and *musculus* (i.e. heterozygosity for ancestry). We counted “junctions” (Fisher 1949; Baird 2006; Janzen et al. 2018) as inferred transitions from one ancestry to another along a chromosome.

With unphased data, we considered the locations of junctions in diploid space, but we could still count junctions in a pseudo-haploid manner. A location where diploid ancestry changes from homozygous (*domesticus* or *musculus*) to heterogenic requires the presence of one junction on one of the two chromosomes. A location where diploid ancestry changes from homozygous *domesticus* to homozygous *musculus* (or the reverse) requires two junctions, one on each chromosome. To be conservative, we required that junctions be identified across analyses using all four pairs of reference populations and we ignored junctions supported by ancestry at single SNPs.

### Reconstruction of Island Colonization

#### THE TWO-POPULATION SITE FREQUENCY SPECTRUM

Our first approach to reconstructing island colonization focused on analyses of the folded, two-dimensional (2D) site frequency spectrum (SFS) for each pair of populations. We did not pursue three-population models because the 3D SFS was too sparse with our sample sizes after close relatives were removed. Treating four mice from Norway with high proportions (>0.9) of *domesticus* ancestry as a mainland reference population, we separately analyzed the 2D SFS for Norway-Sandoy, Norway-Nólsoy, and Sandoy-Nólsoy. We considered putatively neutral SNPs found in regions with the following characteristics: (1) shared *domesticus* ancestry across the three populations, (2) at least 25 kb away from genes, and (3) recombination rates higher than the genomic average (> 0.5 cM/Mb; Cox et al. 2009). Windows were chosen to be at least 5 kb from each other. SNPs were taken from windows ranging from 500 bp to 10 kb in size. The summed genomic length considered was 15,312,841 bp. For each population pair, the 2D SFS was separately generated twice, using two independent approaches: either (1) GATK-called genotypes or (2) ANGSD (Korneliussen et al. 2014) with options ‘-minMapQ 30 -minQ 30 -baq 1’.

To estimate demographic parameters from the 2D SFS, we used the maximum likelihood framework implemented in Diffusion Approximation for Demographic Inference (*dadi*; Gutenkunst et al. 2009) version 2.2.0. This approach employs a diffusion approximation to compute the expected SFS, assuming a specified model of population history. We considered a series of two-population models motivated by island colonization. For mainland-island analyses (Norway-Sandoy and Norway-Nólsoy), potential model parameters included an ancestral effective population size (*N_e_*), an instantaneous change in the ancestral *N_e_*, divergence of the island population from the mainland accompanied by an instantaneous change in island *N_e_*, a constant mainland *N_e_*, an additional instantaneous change in island *N_e_* following its split for the mainland, and migration between the island and the mainland. Population splits and size changes were each associated with an additional parameter that reflects the timing of the event. For the island-island analyses (Sandoy-Nólsoy), the framework was similar, except both populations split directly from an ancestral population and could undergo an additional size change following their split. For all models, we allowed for historical inbreeding in both island mainland populations (Blischak et al. 2020).

To measure model fit and estimate parameters, we ran many replicate analyses for each model/dataset combination. The set of starting parameter values for each replicate was drawn randomly from log-uniform prior distributions with the following ranges (in units of the *dadi* reference population size, which defaults to 1; times are given in units of 2*N_e,ref_* generations, whereas migration rates are in units of 2*N_e,ref_m_ij_*): 10^-4^ to 10 for *N_e_*; 10^-5^ to 10 for event times; 10^-6^ to 20 for migration rates, and 10^-7^ to 1 for inbreeding coefficients. We initially ran 1,000 replicates using parameter bounds recommended by the *dadi* manual. Resulting likelihoods suggested that some bounds should be expanded. We subsequently ran 10,000 replicates using the following bounds: 10^-5^-20 for population sizes; 10^-6^-10 for event times; 0-100 for migration rates; and 0-1 for inbreeding coefficients. If any of the parameter estimates for the top five replicates with highest likelihoods were not in the same order of magnitude, more replicates were run until all parameter estimates converged in order of magnitude. Inferences were based on results from these 10,000 or more replicates, with the best-fit set of parameter values taken to be those in the replicate with the highest log-likelihood. Parameter estimates were rescaled to units of interest (*e.g.* time in generations) following recommendations in the *dadi* manual. To identify best-fit models, we compared log-likelihoods among nested models through adjusted likelihood ratio tests (Coffman et al. 2016). Likelihood ratio tests comparing all models evaluated are available in **Table S7**. We also visually compared plots of residuals between observed and expected 2D SFS among all models. Parameter uncertainty for best-fit models was quantified using the Godambe Information Matrix (Coffman et al. 2016) with 200 bootstrap replicates.

Recognizing the potential for uncertainty in reconstructing the SFS, we performed separate demographic analyses that used either GATK-derived 2D SFS or ANGSD-derived 2D SFS as input datasets.

#### THE ANCESTRAL RECOMBINATION GRAPH

Our second approach to reconstructing island colonization focused on the ancestral recombination graph (ARG). We used the ARG-based method *Relate* v1.2.2 (Speidel et al. 2019) to infer historical changes in *N_e_* and to estimate the split times between populations. *Relate* infers topological features of local genealogies based on position-specific distance matrices between haplotypes. Following the reconstruction of local tree topology, *Relate* estimates branch lengths using a Markov chain Monte Carlo algorithm, initially assuming constant *N_e_*. Estimated branch lengths are then used to re-estimate the historical trajectory of *N_e_* as the inverse of the average coalescence rate during each epoch, enabling the re-estimation of branch lengths under the inferred demography. This process is iteratively repeated to co-estimate branch lengths and the trajectory of *N_e_*.

For this set of analyses, we included all sequences from unrelated mice from Nólsoy, Sandoy, and Norway, regardless of inferred subspecific ancestry. To make inferences from the ARG more directly comparable to inferences from the SFS, we conducted a separate ARG analysis for each pair of populations: Norway-Sandoy, Norway-Nólsoy, and Sandoy-Nólsoy. *Relate* requires that SNP alleles be polarized as ancestral or derived and haplotypes be phased. To polarize GATK-filtered SNPs we examined genotype calls from genome sequences of two outgroups, *Mus caroli* (Thybert et al. 2018) and *Mus spretus* (Keane et al. 2011), aligned to the mouse reference genome sequence. For each species, we downloaded conversion files between genome assemblies from the UCSC Genome Browser and used UCSC ‘LiftOver’ to find the positions of variants. To be conservative, we required that these two outgroups harbored the same allele, which we inferred to be the ancestral allele. SNPs that did not meet this criterion were eliminated. Haplotypes were phased separately for each population by passing polarized SNP calls to *Beagle* v5.4 (Browning et al. 2021). We assigned to SNPs positions on the genetic map using linear interpolation from the mouse reference genetic map (Cox et al. 2009). Phased variant call format (VCF) files were merged and converted to the hap/sample format using RelateFileFormats –mode ConvertFromVcf. The haps, sample, and genetic map files were passed to *Relate* v1.2.2 (Speidel et al. 2019) to perform genome-wide genealogy estimation assuming a mutation rate of 5 x 10^-9^ (per-site, per-generation) and an initial *N_e_* of 20,000. Historical *N_e_* values were calculated from estimated coalescence rates as ½ divided by the coalescence rate. Coalescence rates were calculated from the genealogy results using the ‘EstimatePopulationSize’ module in *Relate* with ‘year_per_generation’ = 0.5. Population split times were inferred as the times when the inverse of the cross-population coalescence rates were the smallest (Kingman 1982). One hundred replicates initiated with different seed numbers were collected to assess convergence.

### Genomic Scans for Selection

To locate genomic regions with evidence for positive directional selection in Faroe Island mice, we used three strategies. First, we searched for genomic regions showing a skew toward rare alleles in the single-population SFS or an elevation in differentiation between mainland-island pairs of populations that were unusual under the inferred demographic history. We restricted these analyses to genomic regions with shared *domesticus* ancestry (again excluding those masked for repeats and segmental duplications). We separated 5kb windows into 4 bins based on the number of accessible (unfiltered) sites: 1-2kb, 2-3kb, 3-4kb, and 4-5kb. Windows smaller than 1kb were discarded. We used scikit-allel v1.3.6 (https://github.com/cggh/scikit-allel) to calculate summary statistics of variation. We computed nucleotide diversity (Tajima 1983) and Tajima’s D (Tajima 1989) separately for Nólsoy, Sandoy, and Norway in these windows. We computed F_ST_ (Hudson et al. 1992) for Norway-Sandoy and Norway-Nólsoy in these windows. For each bin and summary statistic, we counted the number of windows with non-NA values and non-infinite values. To reconstruct the null distribution of Tajima’s D and F_ST_ expected under neutrality, we used *msprime* (Baumdicker et al. 2022) to conduct neutral simulations under the demographic models that provided the best fits to the GATK-derived SFS (from *dadi*) without inbreeding. We assumed the mutation rate and recombination rate were 5 x 10^-9^ per site per generation. We simulated ten times the number of windows, separately for each bin and summary statistic. We treated the Tajima’s D values and F_ST_ values resulting from these simulations as null distributions for genomic scans for selection. For each window in the observed dataset, we computed a *p*-value for Tajima’s D as the proportion of simulated windows with values more negative than the observed value. For each window in the observed dataset, we computed a *p*-value for F_ST_ as the proportion of simulated windows with values higher than the observed value. Windows with *p*-values less than 0.05 divided by the number of tested windows were declared significant after a Bonferroni correction. This conservative approach accounts for the characteristics of the dataset, demographic history, and the performance of many tests along the genome.

In a second strategy to search the genome for positive selection, we compared the local SFS to the genomic SFS, using the method implemented in *Sweepfinder2* (De Giorgio et al. 2016). For this analysis, we considered SNPs in regions of shared *domesticus* ancestry (again excluding those masked for repeats and segmental duplications). Nólsoy, Sandoy, and Norway were analyzed separately using the same grids with 20 kb spacing to enable direct comparisons. The background SFS was computed using all autosomal SNPs for each population. Recombination rates were again estimated by linear extrapolation from the mouse reference genetic map (Cox et al. 2009). We used the following *SweepFinder2* command: ‘SweepFinder2 - lru grids_chrNum.txt popName_chrNum_sf2_alleleFreq.txt popName_SpectFile popName_chrNum_sf2_recMap.txt sf2_grids_output_popName_chrNum.txt’. For each window, *SweepFinder2* computes a likelihood ratio test that compares the fit of a model allowing the local SFS to differ from the background SFS to the fit of a model assuming the local SFS is drawn from the background SFS. To reconstruct null distributions of the likelihood ratio test statistic, we used *msprime* to separately simulate neutral datasets for each pair of mainland and island populations (Norway-Sandoy and Norway-Nólsoy) following the demographic models that provided the best fits to the GATK-derived SFS without inbreeding. We simulated 20,000 replicates of a 1 Mb sequence with mutation rate and recombination rate set to 5 x 10^-9^ per site per generation. The background SFS was reconstructed from all 20,000 replicates. We conducted a *SweepFinder2* scan on each of the 20,000 simulated datasets, using a grid size of 20 kb with windows centered at the grid points and treating the full SFS across all simulations as the background SFS. For each 20 kb window in the observed dataset, we compared the likelihood ratio test statistic to the simulated distribution of this statistic to compute a *p*-value under the null hypothesis of neutrality. Windows with *p*-values less than 0.05 divided by the number of tested windows were declared significant after a Bonferroni correction. This conservative approach again accounts for the characteristics of the dataset, demographic history, and the performance of many tests along the genome.

Our third tactic for scanning the genome for positive selection focused on the ARG. For this analysis, we separately considered Nólsoy, Sandoy, and Norway, using the set of polarized, GATK-filtered SNPs that were used in the demographic reconstruction by *Relate*. The set of variants included in this ARG analysis was much larger than the sets of variants included in the other two genomic scan approaches, since it did not require that we filter for subspecific ancestry. *Relate* evaluates the evidence for selection at a SNP by estimating how quickly the lineage carrying the mutation spreads through the population relative to other lineages (Speidel et al. 2019). The program treats the number of present-day carriers of the mutation as a test statistic, conditioning on the number of lineages present when the mutation first enters the population. The null distribution of the test statistic is computed analytically and is claimed to be robust to changes in population size (Speidel et al. 2019). Our selection scans followed recommendations in the *Relate* user’s manual. Population size history was estimated using the setting –threshold 0, so that the branch lengths of all trees were updated for the estimated population size history. SNPs with low derived allele frequencies were filtered out using RelateFileFormats –mode GenerateSNPAnnotations to annotate .mut files. *Relate* provides a *p*- value for each SNP under the null hypothesis of neutrality. We used a Bonferroni adjustment to account for multiple testing, considering the number of SNPs or the estimated number of trees as the number of tests in separate corrections. To explore evidence for parallel evolution across island populations, we identified the subset of SNPs with uncorrected *P* < 0.05 in both Nólsoy and Sandoy. To search for enrichment of functions among genes in which these SNPs were located, we conducted a gene ontology analysis using g:Profiler (https://biit.cs.ut.ee/gprofiler/gost).

## Data Availability

Code used for the main analyses is available on GitHub (https://github.com/PayseurLabUWMadison/Faroe_Demography). Upon submission, raw sequencing reads will be deposited in the SRA and VCF files containing subspecific ancestry calls and genotype calls used for analyses of demographic history will be uploaded to Dryad.

## Acknowledgments

We thank the following individuals for assistance with field collections: Emilie Hardouin and the Johannessens on Mykines, Noomi Gregersen and Heidi S. Hansen in the Faroe Islands, and Jeroen van der Kooij in Norway. We thank Mark Nolte for assistance with DNA extractions. Members of the Payseur lab provided helpful input during the project.

## Author Contributions

B.A.P. designed the study, with assistance from P.J., M.E.F., and E.K.H. E.P.J., E.M., J.- K.M., Y.F.C., and J.B.S. acquired samples used for the study. P.J. conducted the research, with contributions from M.E.F. and E.K.H. and supervision from B.A.P. B.A.P. wrote the manuscript, with substantive input from E.K.H., P.J., J.B.S., and M.E.F.

