## Supplemental Figures S1-S4 for "Population Genomics of Giant Mice from the Faroe Islands: Hybridization, Colonization, and a Novel Challenge to Identifying Genomic Targets of Selection"

**Figure S1. Subspecies ancestries of mice from the Faroe Islands and Norway.** Each page depicts one chromosome. Each row is one mouse. Subspecies ancestries are color-coded.

Figure S1

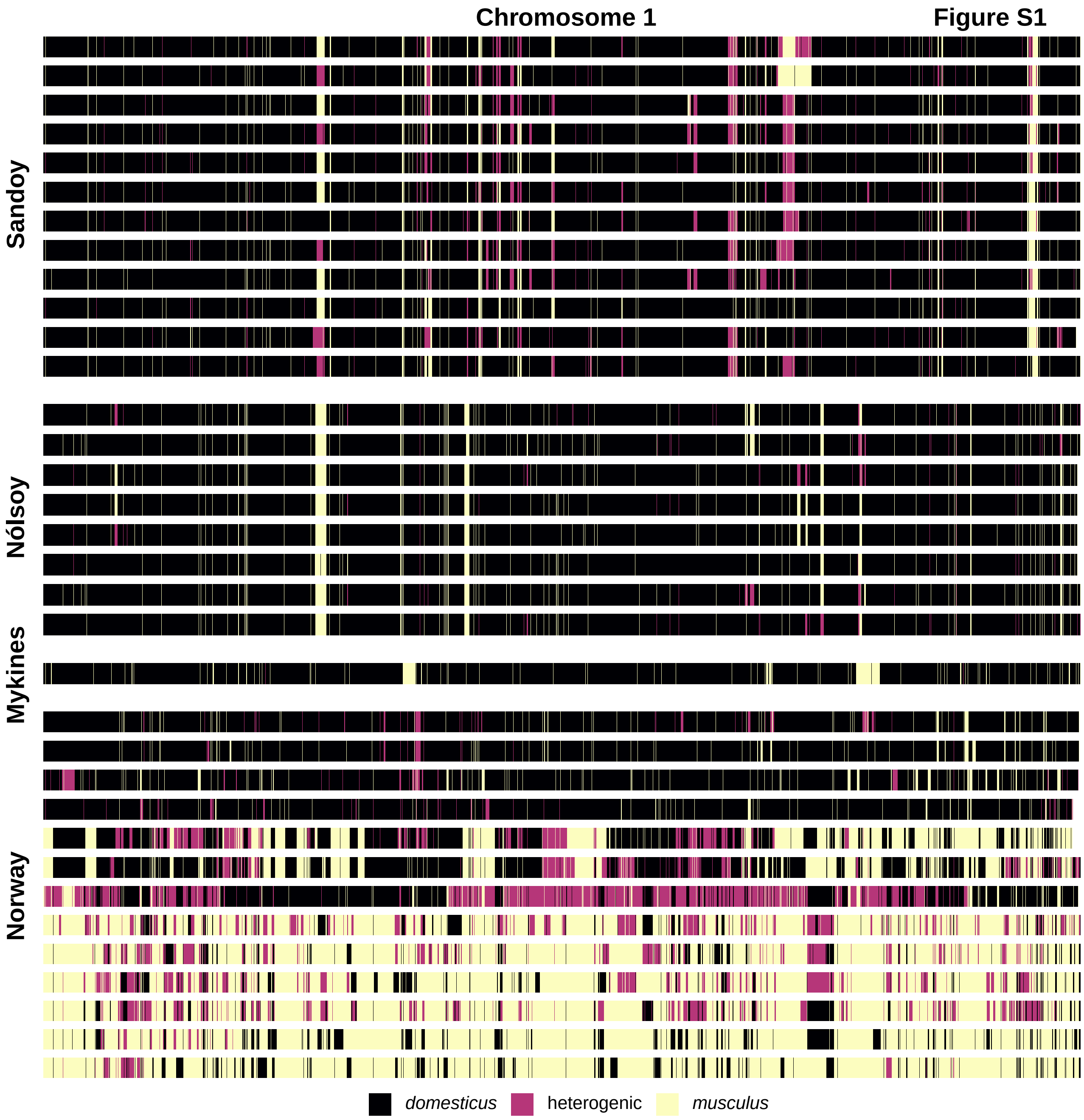
Chromosome 1
Sandoy
Nólsoy
Mykines
Norway
domesticus
heterogenic
musculus

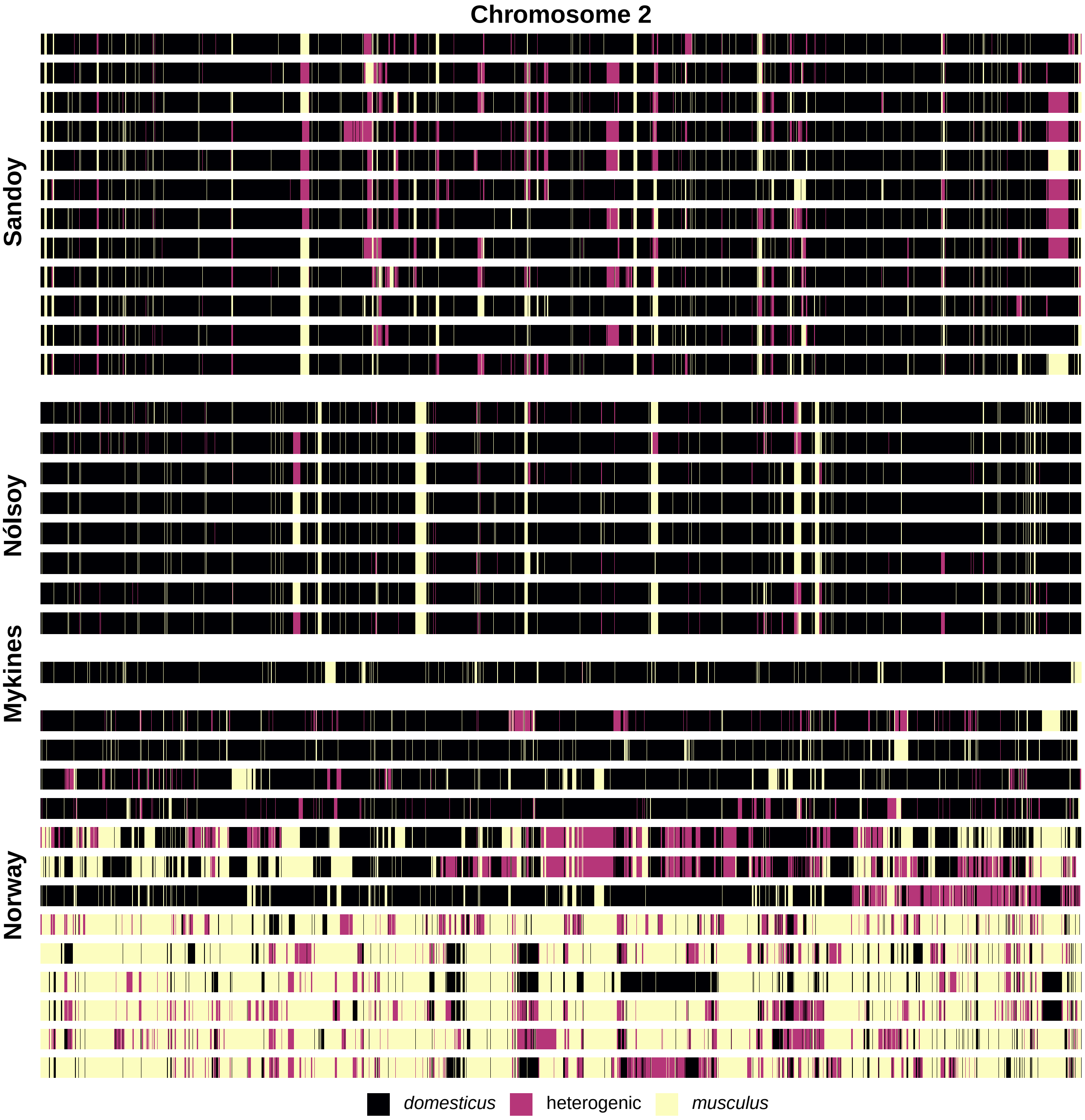
Chromosome 2
Sandoy
Nólsoy
Mykines
Norway
domesticus
heterogenic
musculus

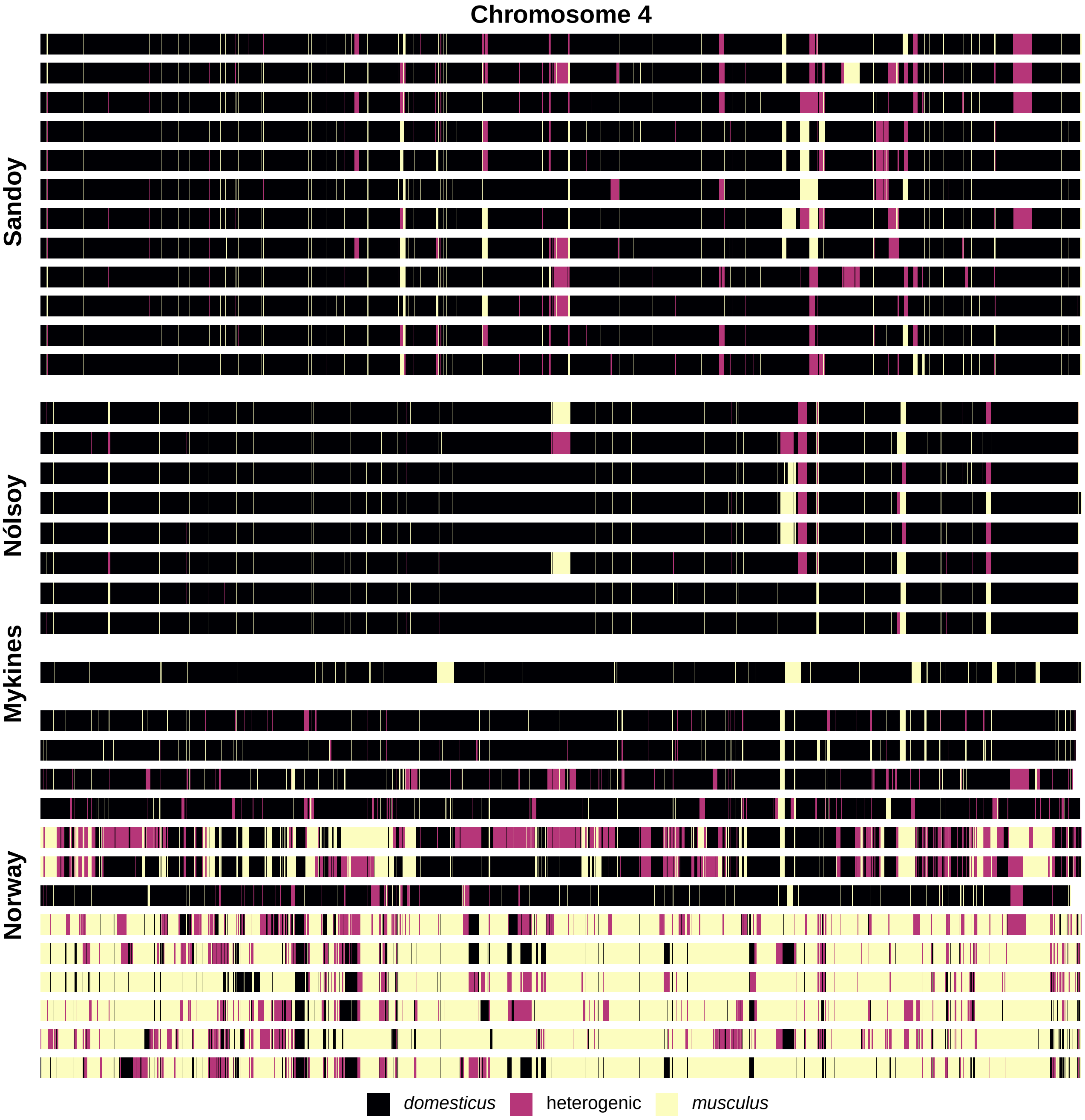

Chromosome 4
Sandoy
Nólsoy
Mykines
Norway
domesticus
heterogenic
musculus

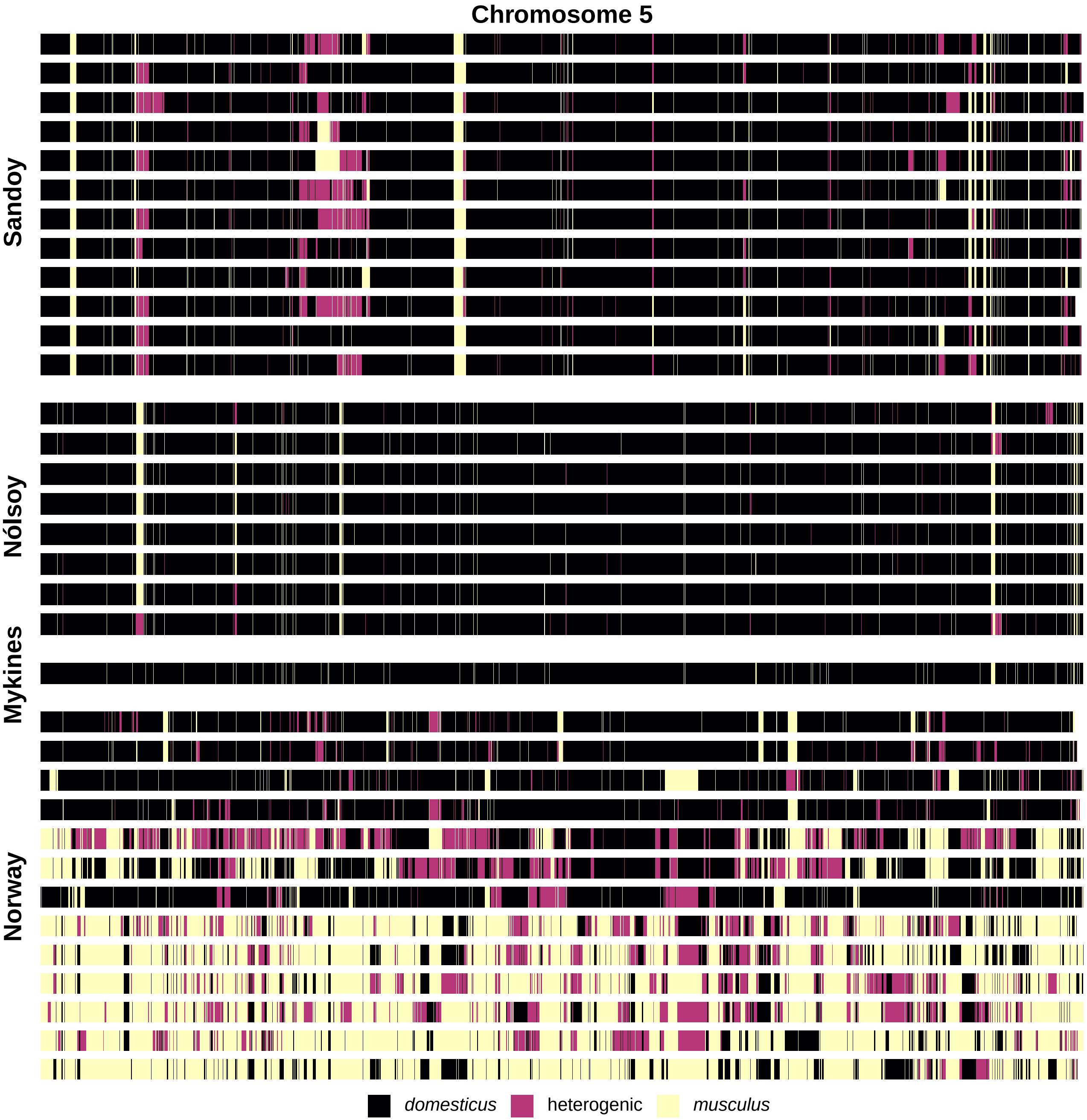

Chromosome 5
Sandoy
Nólsoy
Mykines
Norway
domesticus
heterogenic
musculus

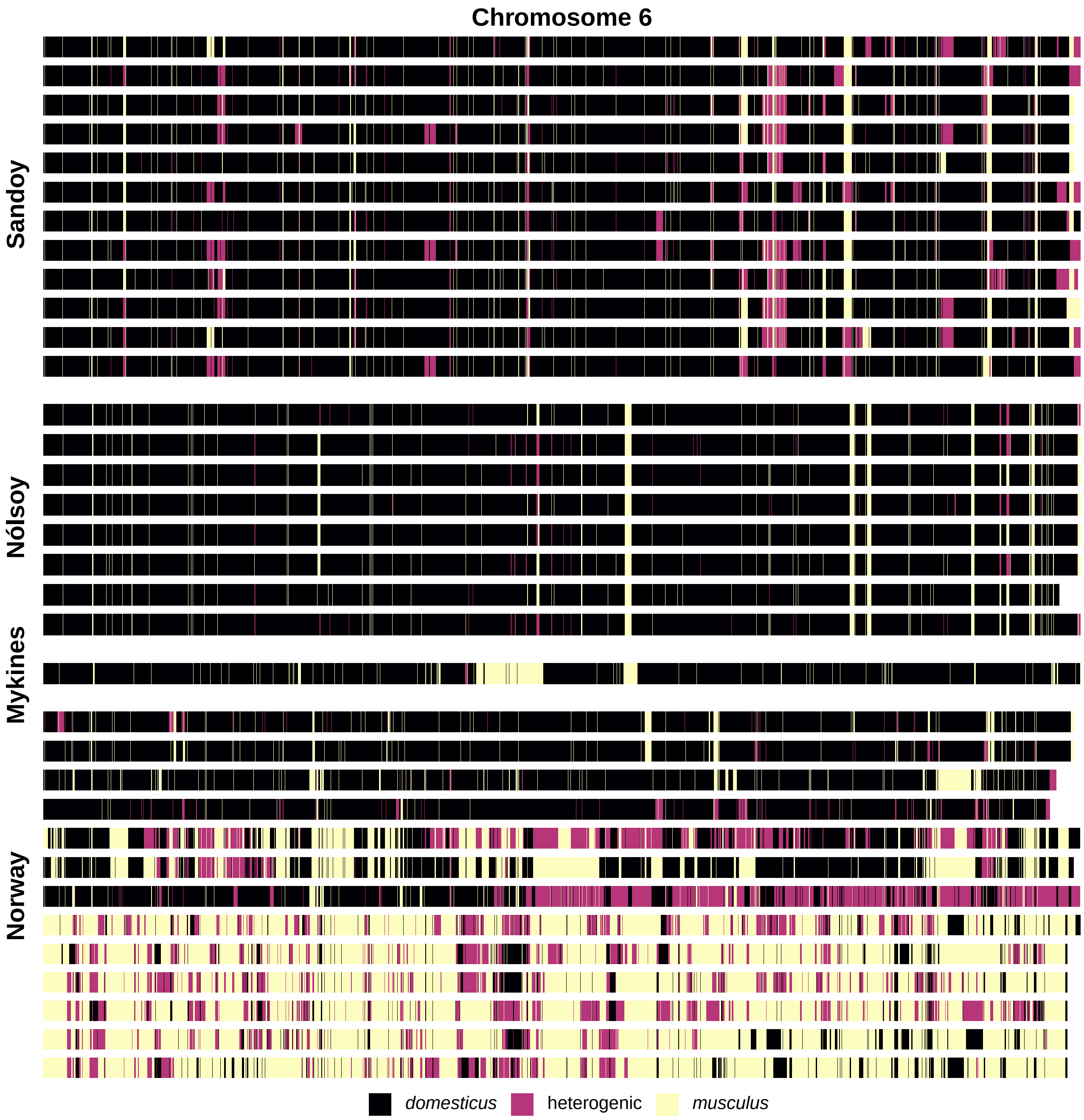
Chromosome 6
Sandoy
Nólsoy
Mykines
Norway
domesticus
heterogenic
musculus

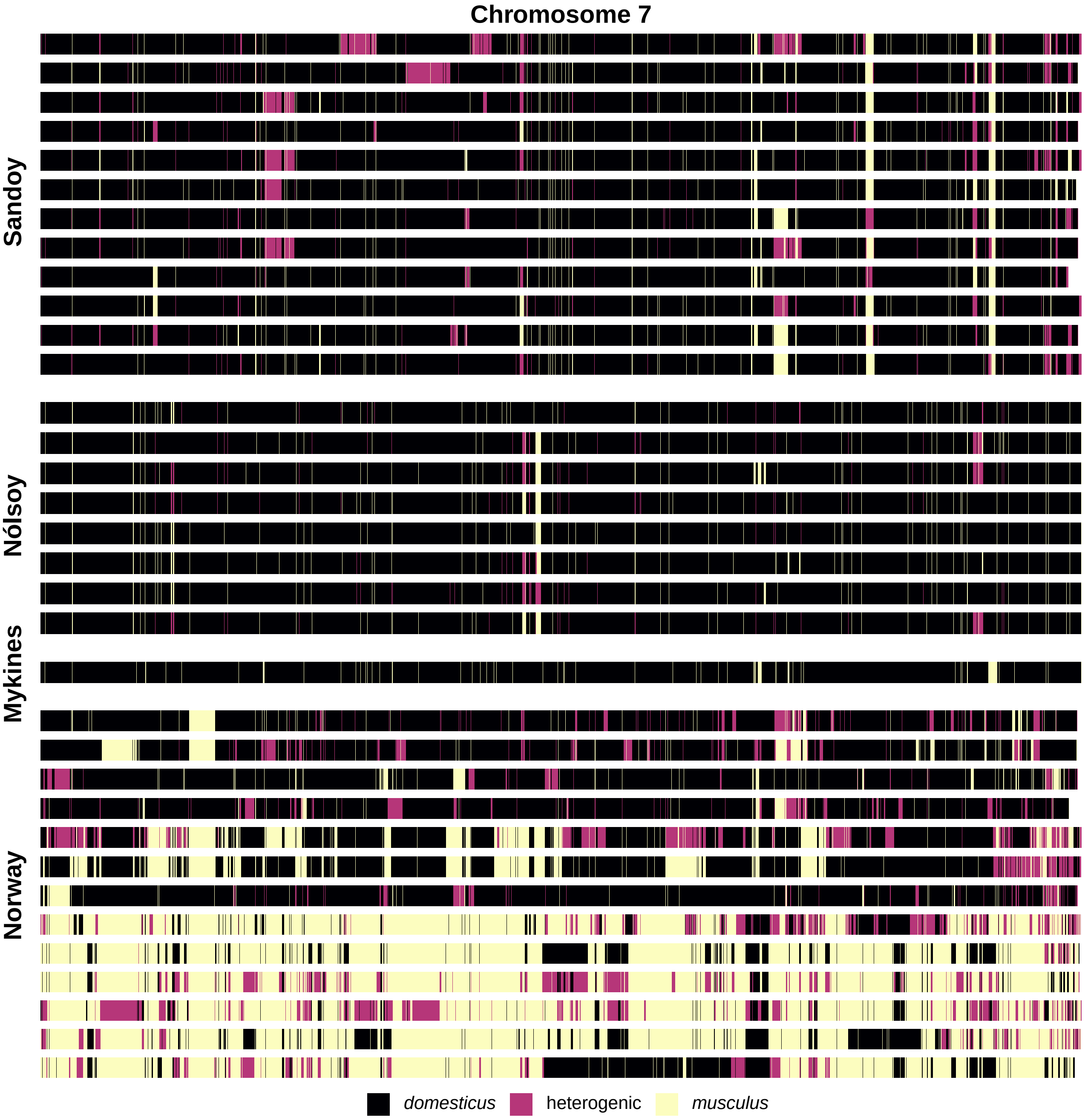
Chromosome 7
Sandoy
Nólsoy
Mykines
Norway
domesticus
heterogenic
musculus

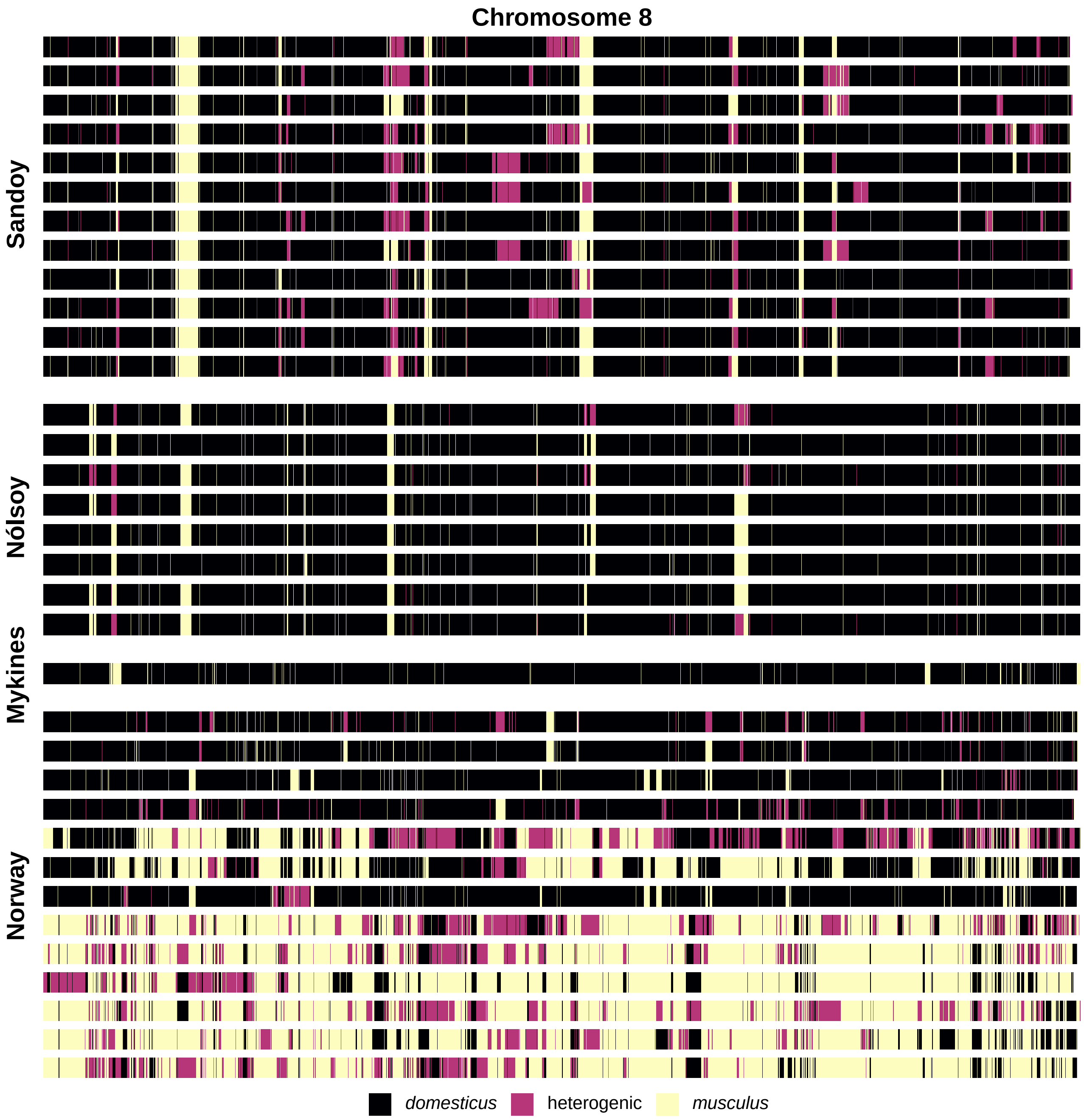
Chromosome 8
Sandoy
Nólsoy
Mykines
Norway
domesticus
heterogenic
musculus

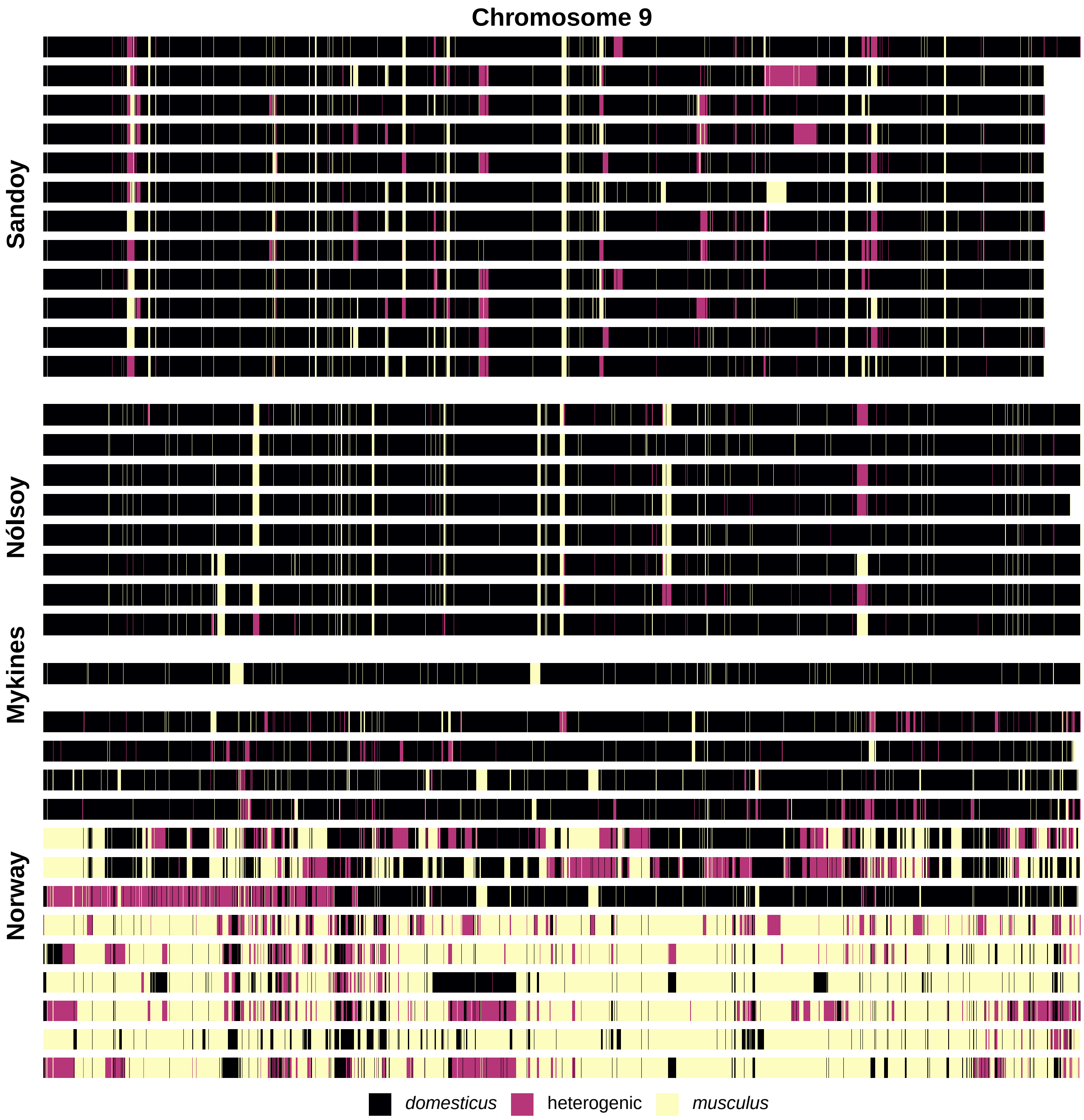
Chromosome 9
Sandoy
Nólsoy
Mykines
Norway
domesticus
heterogenic
musculus

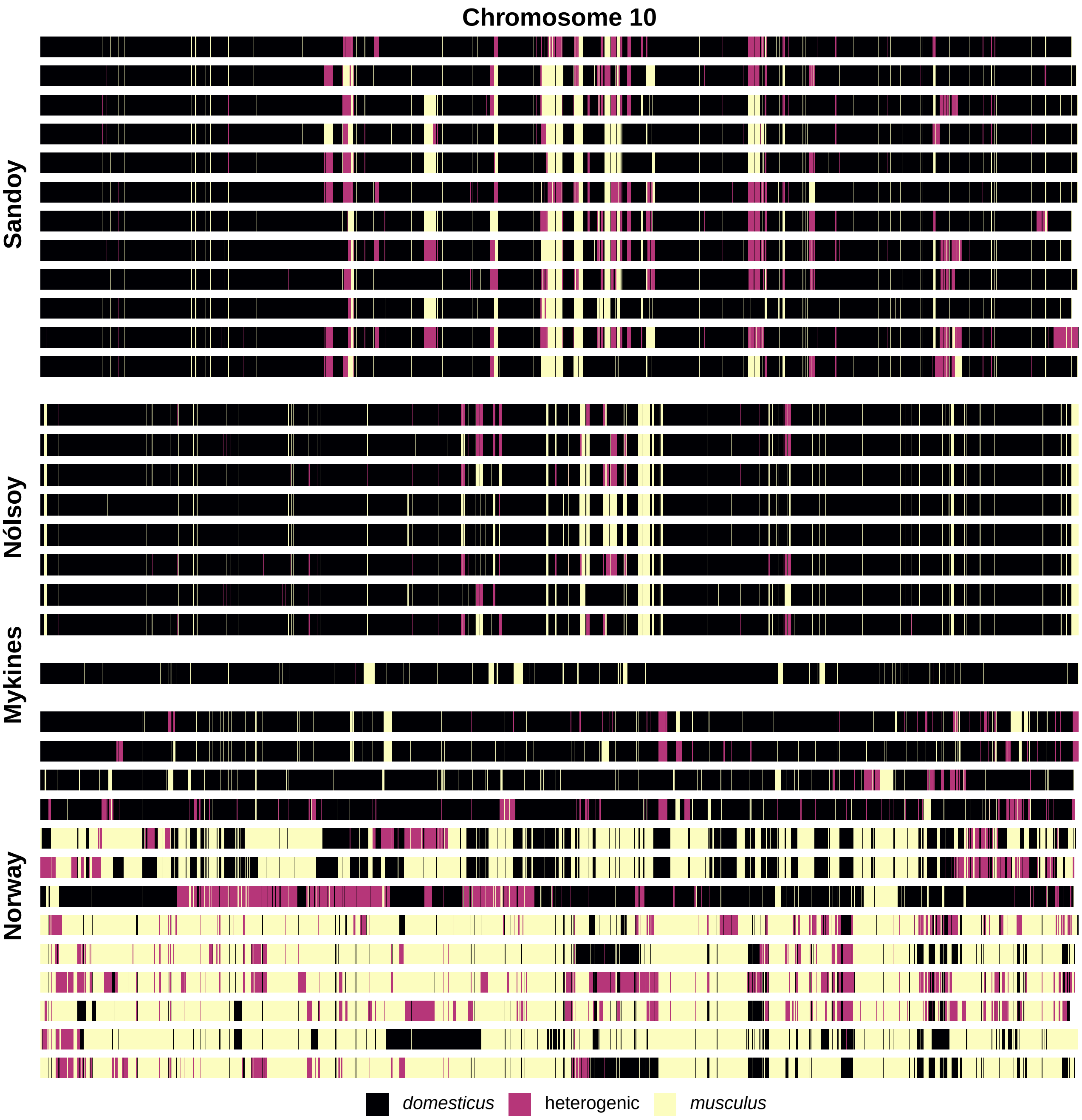

Chromosome 10
Sandoy
Nólsoy
Mykines
Norway
domesticus
heterogenic
musculus

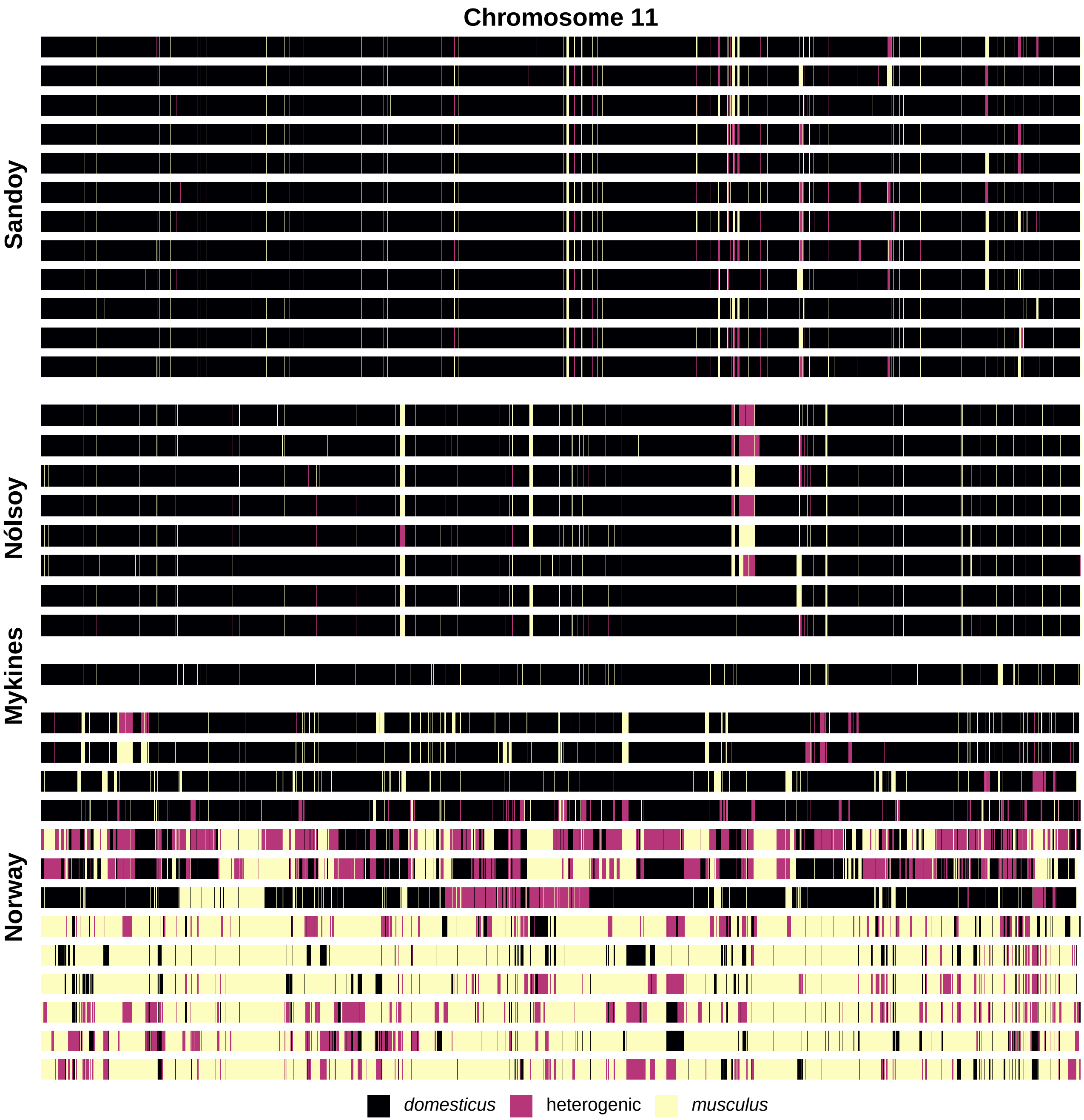
Chromosome 11
Sandoy
Nólsoy
Mykines
Norway
domesticus
heterogenic
musculus

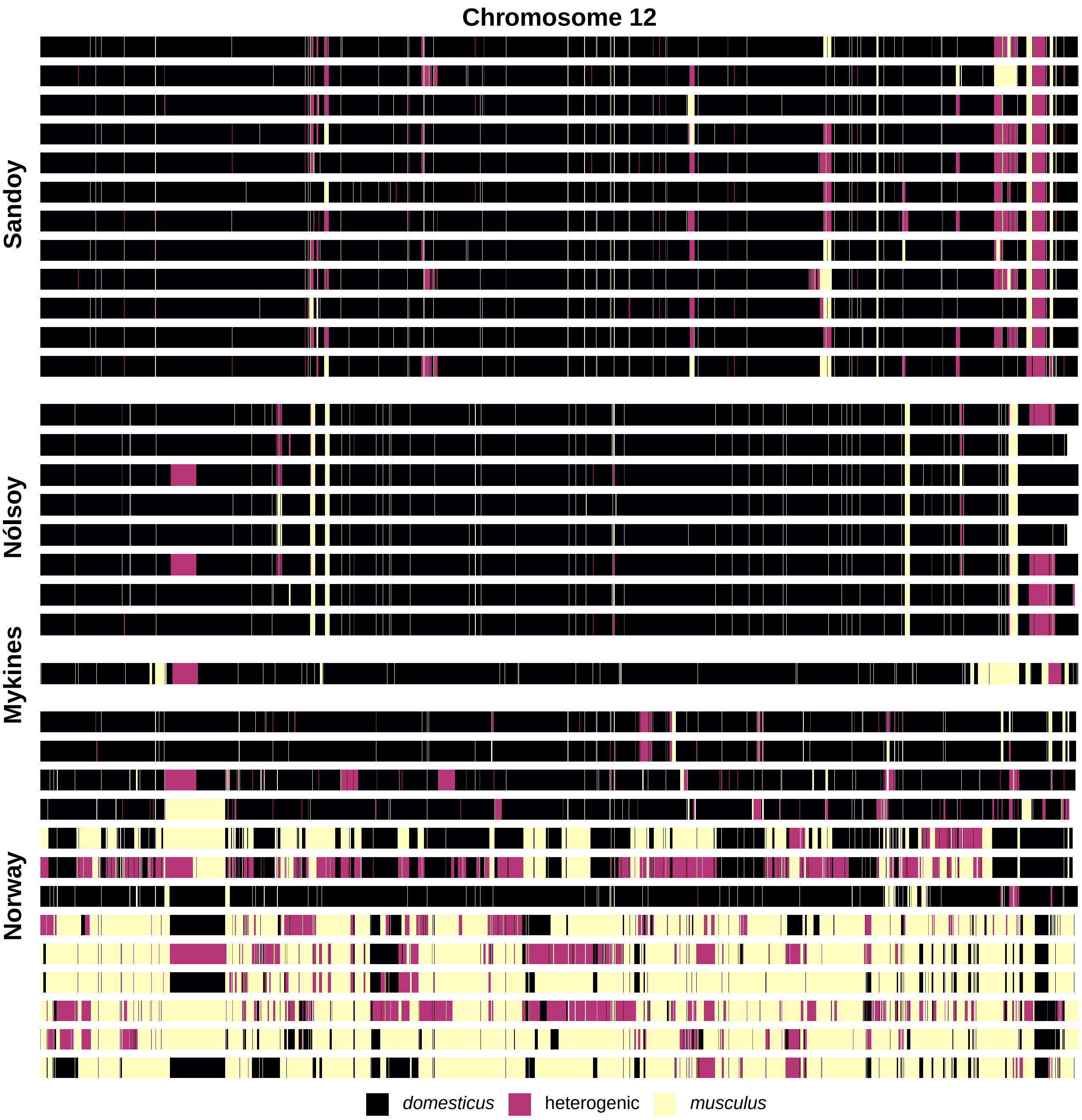
Chromosome 12
Sandoy
Nólsoy
Mykines
Norway
domesticus
heterogenic
musculus

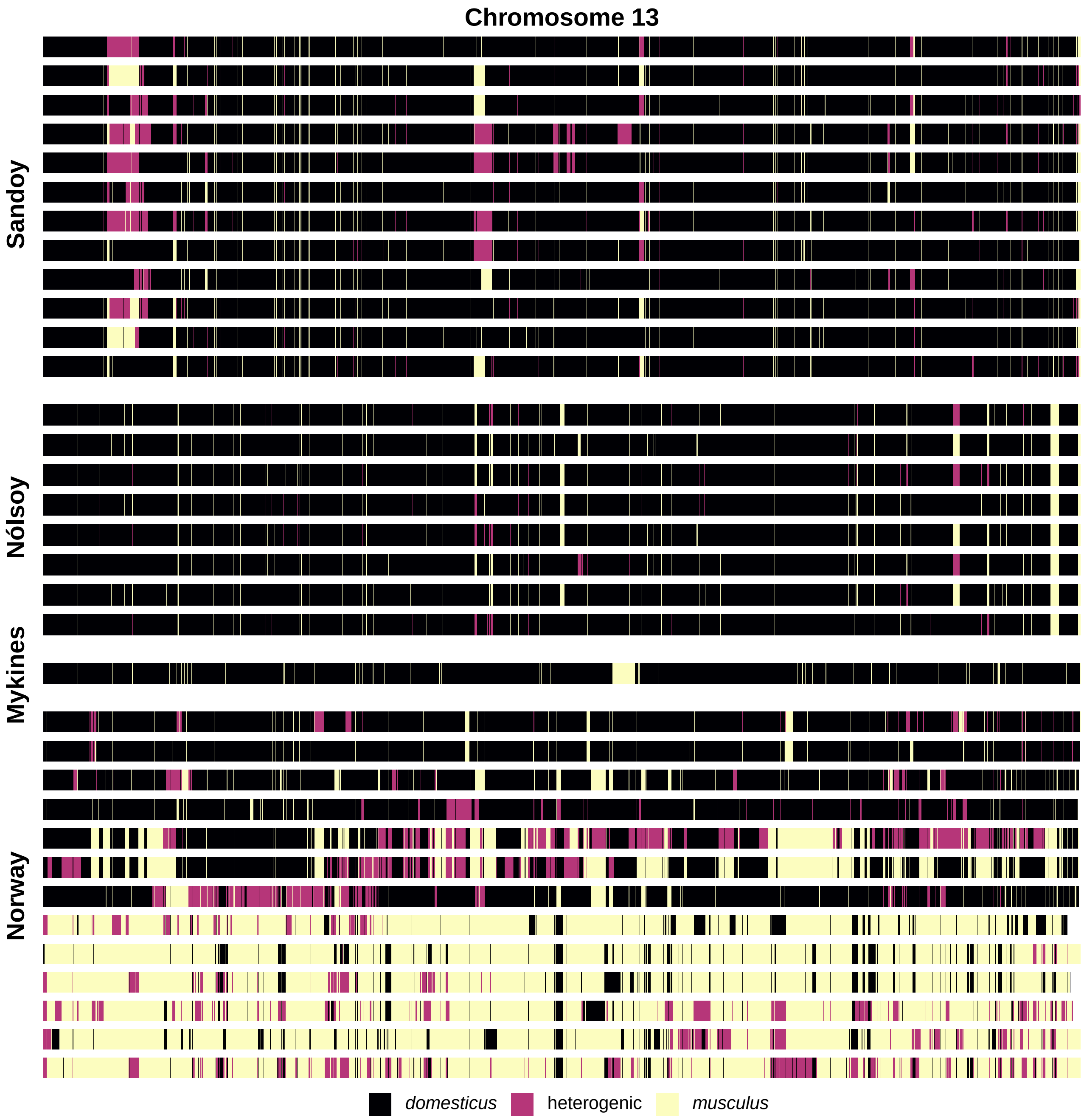
Chromosome 13
Sandoy
Nólsoy
Mykines
Norway
domesticus
heterogenic
musculus

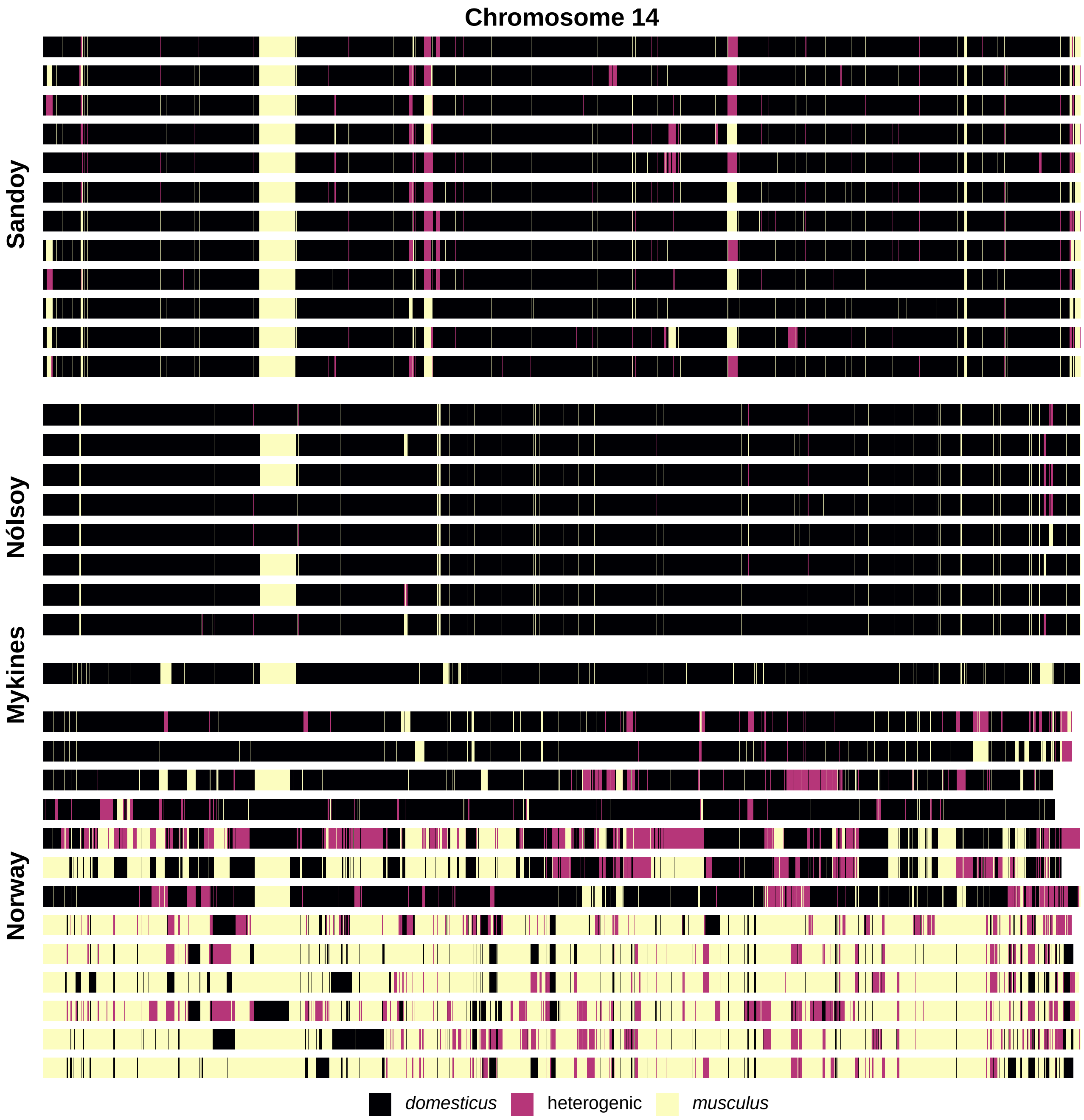
Chromosome 14
Sandoy
Nólsoy
Mykines
Norway
domesticus
heterogenic
musculus

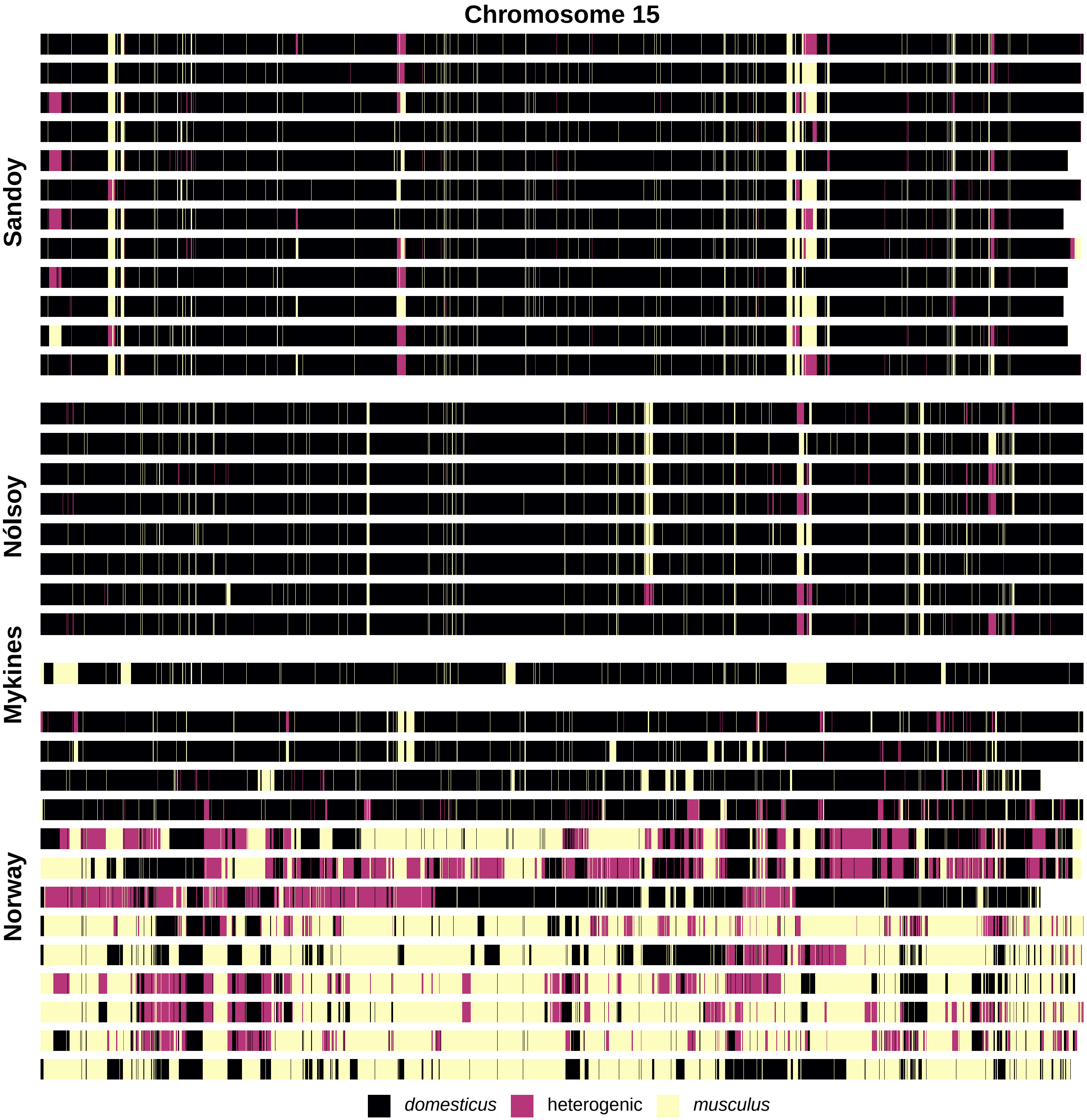
Chromosome 15
Sandoy
Nólsoy
Mykines
Norway
domesticus
heterogenic
musculus

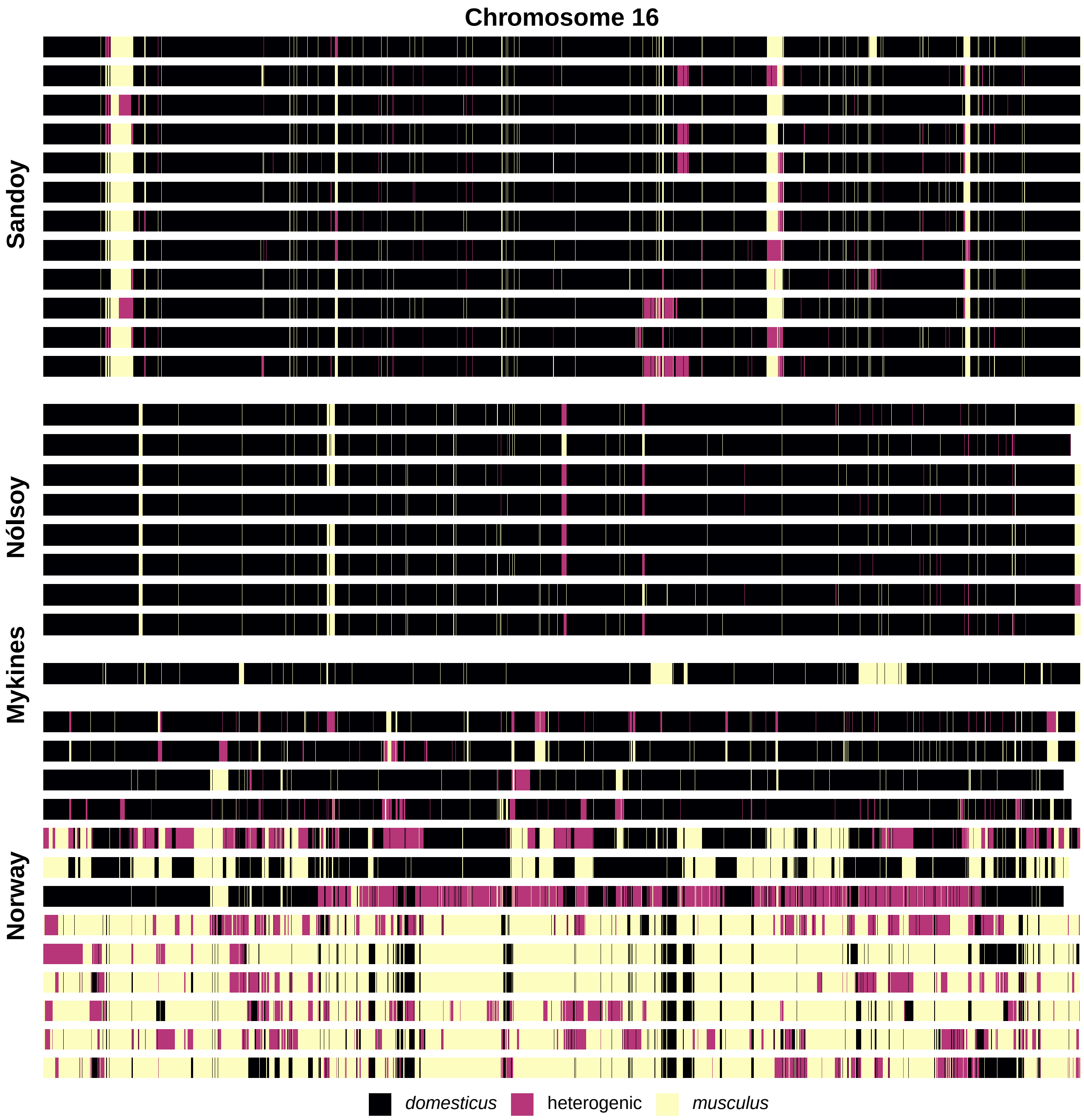
Chromosome 16
Sandoy
Nólsoy
Mykines
Norway
domesticus
heterogenic
musculus

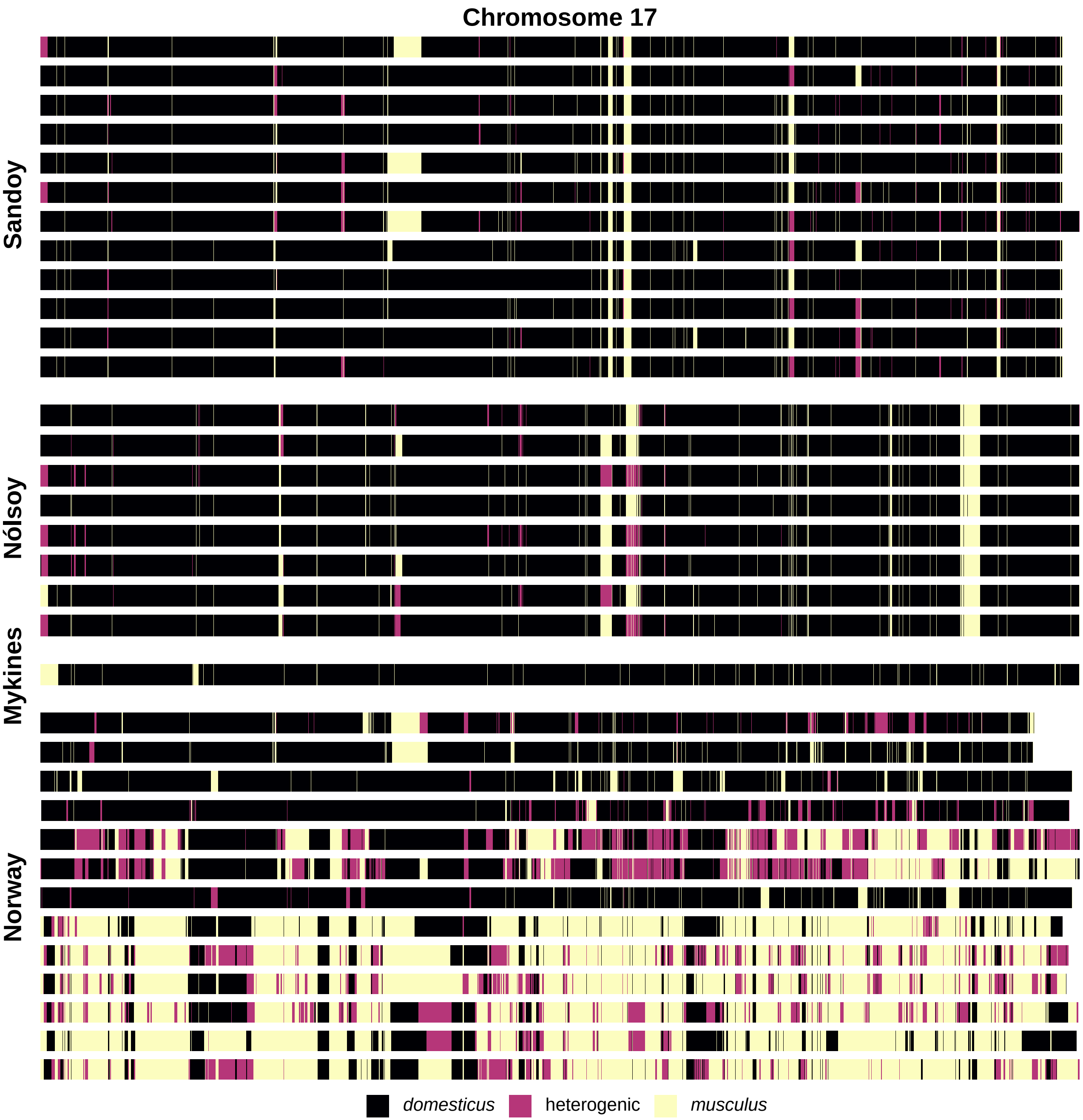

Chromosome 17
Sandoy
Nólsoy
Mykines
Norway
domesticus
heterogenic
musculus

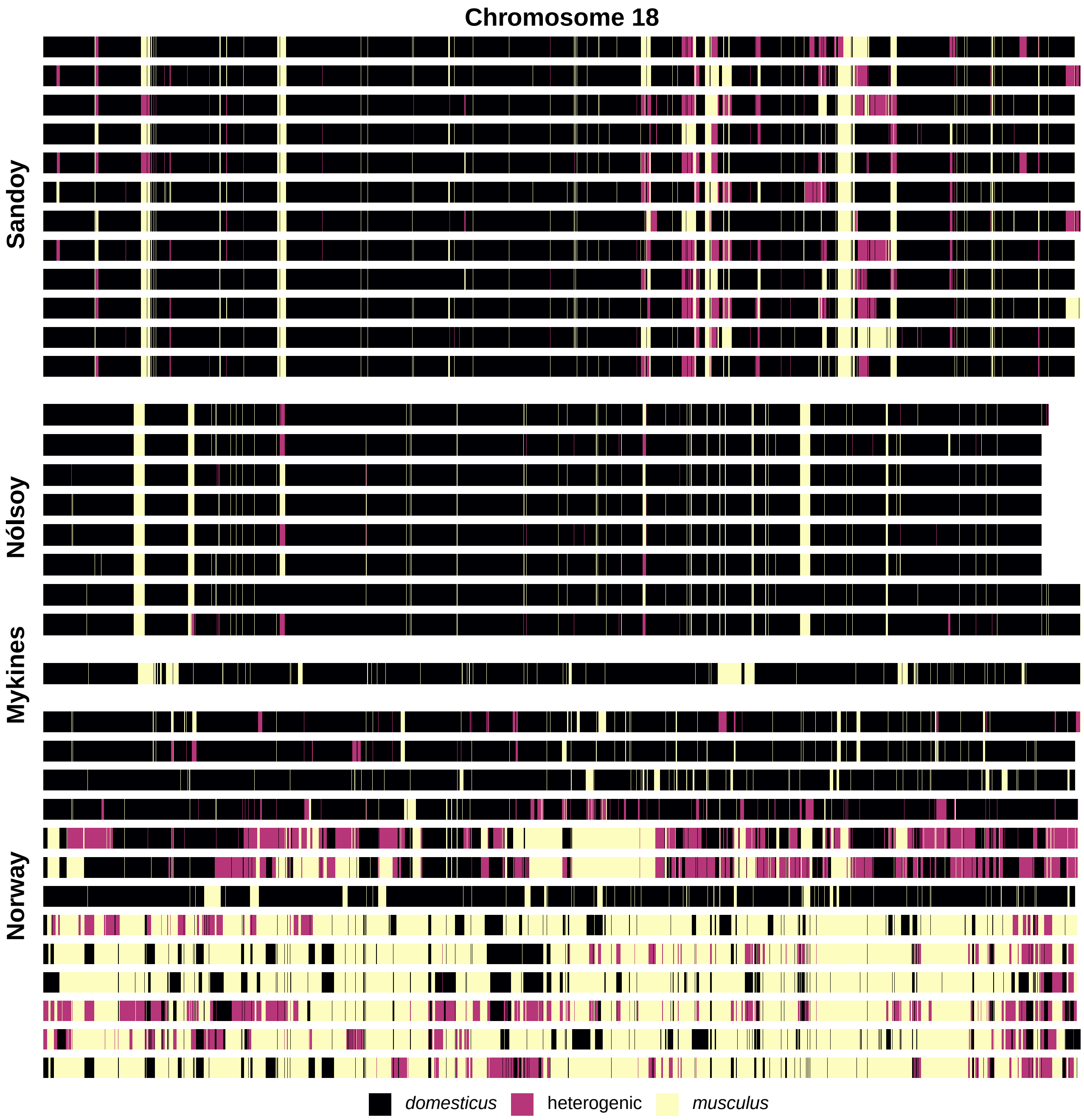

Chromosome 18
Sandoy
Nólsoy
Mykines
Norway
domesticus
heterogenic
musculus

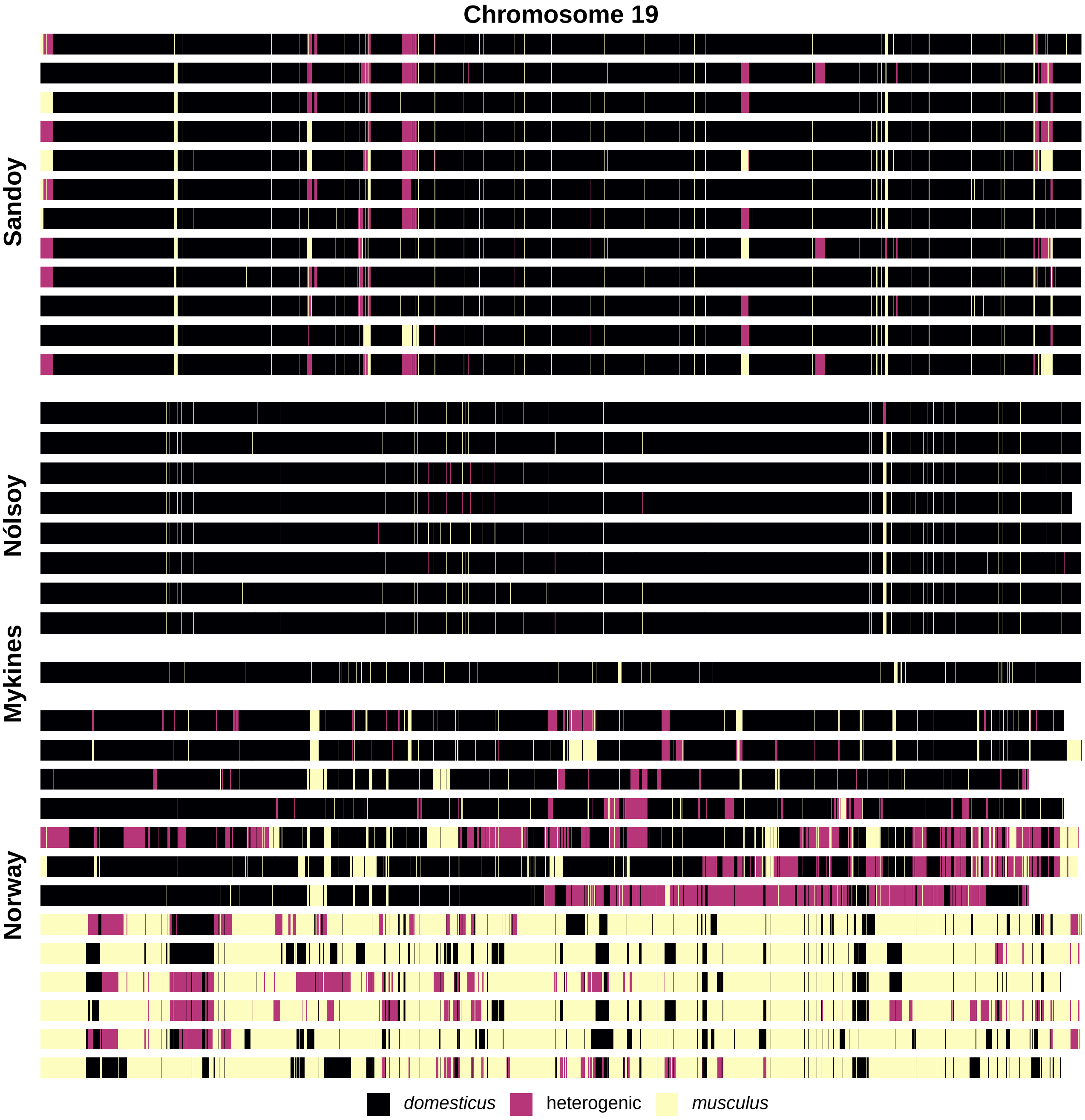
Chromosome 19
Sandoy
Nólsoy
Mykines
Norway
domesticus
heterogenic
musculus

**Figure S2. Best-fitting demographic models for mice from Nólsoy and Sandoy, treating mice from Norway as a mainland reference population.** Parameter estimates come from analyses of two-dimensional site frequency spectra for pairs of populations inferred by ANGSD for putatively neutral SNPs in genomic regions with shared *M. m. domesticus* ancestry. $N_e$ = effective population size. $t$ = timing in generations of population splits or changes in $N_e$, ordered beginning with the most recent event. $m$ = migration rate. Population colors match Figure 1, Figure 3, Figure 4, and Figure S3.

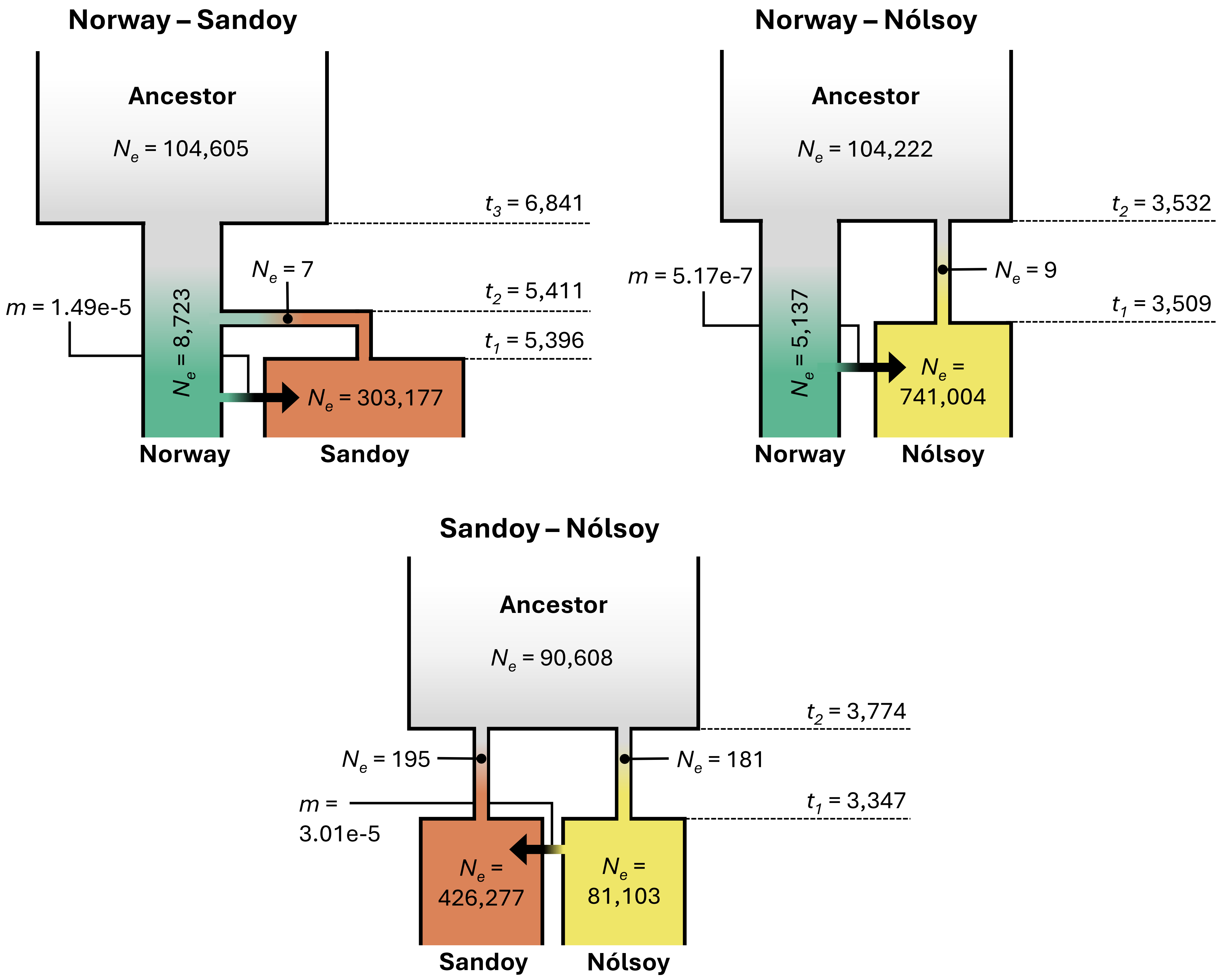
Norway – Sandoy
Ancestor
$N_e$ = 104,605
$t_3$ = 6,841
$N_e$ = 7
$t_2$ = 5,411
$t_1$ = 5,396
$m$ = 1.49e-5
$N_e$ = 8,723
$N_e$ = 303,177
Norway
Sandoy
Norway – Nólsoy
Ancestor
$N_e$ = 104,222
$t_2$ = 3,532
$N_e$ = 9
$t_1$ = 3,509
$m$ = 5.17e-7
$N_e$ = 5,137
$N_e$ = 741,004
Norway
Nólsoy
Sandoy – Nólsoy
Ancestor
$N_e$ = 90,608
$t_2$ = 3,774
$N_e$ = 195
$N_e$ = 181
$t_1$ = 3,347
$m$ = 3.01e-5
$N_e$ = 426,277
$N_e$ = 81,103
Sandoy
Nólsoy

**Figure S3. Effective population sizes reconstructed from ancestral recombination graphs.** Parameter estimates come from analyses of ancestral recombination graphs for pairs of populations based on GATK-derived genotypes, with alleles polarized by comparison to two outgroup species. Each panel shows one cross-population comparison and two within-population comparisons. Population Size = effective population size. Vertical dotted lines denote estimated split times between each pair of populations. Note that the x-axis appears on a logarithmic scale. Colors match Figure 1, Figure 3, Figure 4, and Figure S2.

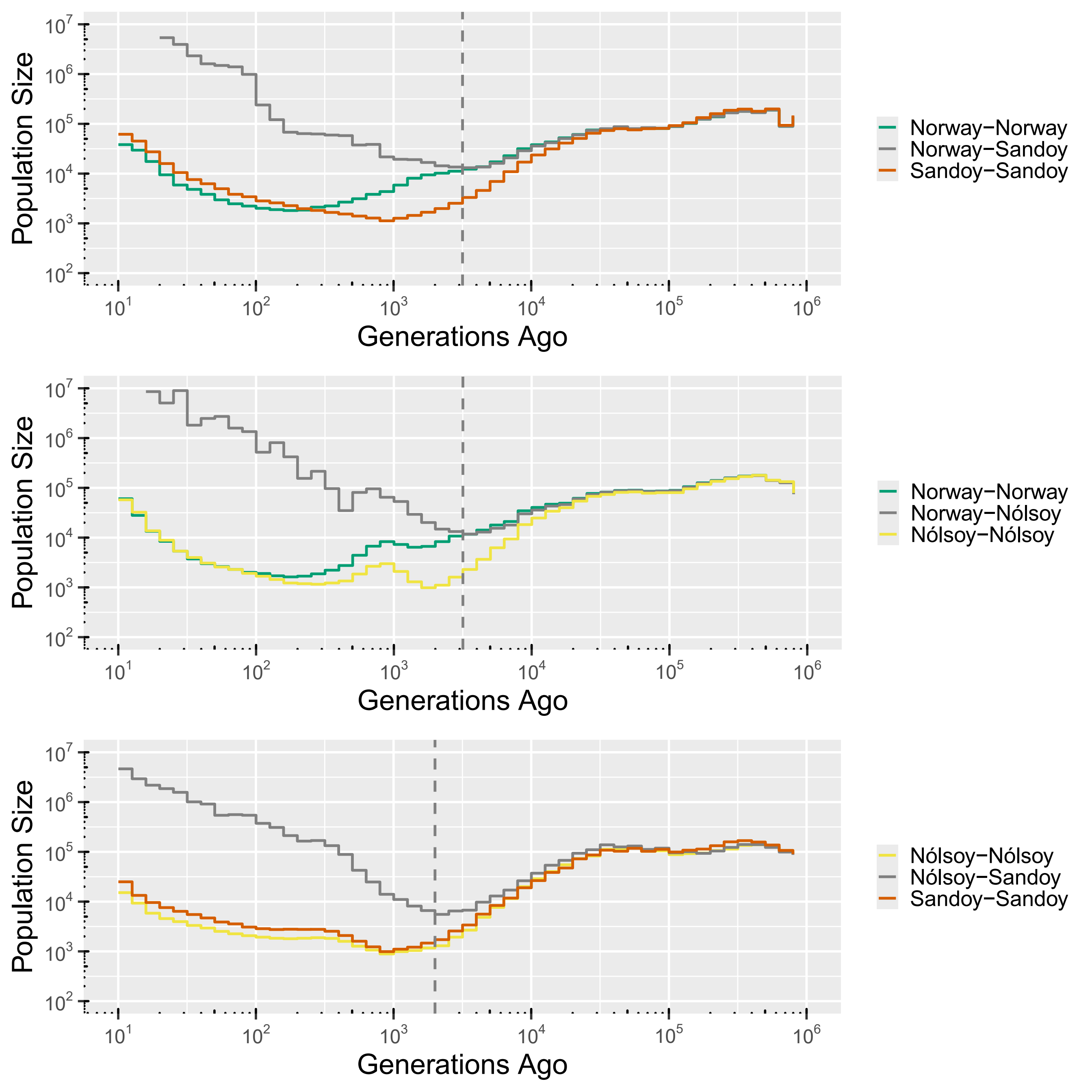
Population Size
Generations Ago
Norway−Norway
Norway−Sandoy
Sandoy−Sandoy
Population Size
Generations Ago
Norway−Norway
Norway−Nólsoy
Nólsoy−Nólsoy
Population Size
Generations Ago
Nólsoy−Nólsoy
Nólsoy−Sandoy
Sandoy−Sandoy

**Figure S4. Distributions of nucleotide diversity, Tajima's D, and Fst for genomic windows compared to simulations.** Simulations followed best-fitting models without inbreeding to the two-dimensional site frequency spectrum inferred from GATK-derived genotypes for putatively neutral SNPs in genomic regions with shared *M. m. domesticus* ancestry. Each page compares distributions of a single summary statistic computed for two populations for windows throughout genomic regions with shared *M. m. domesticus* ancestry and from simulations. Distributions drawn from empirical data are labeled by population.

Figure S4

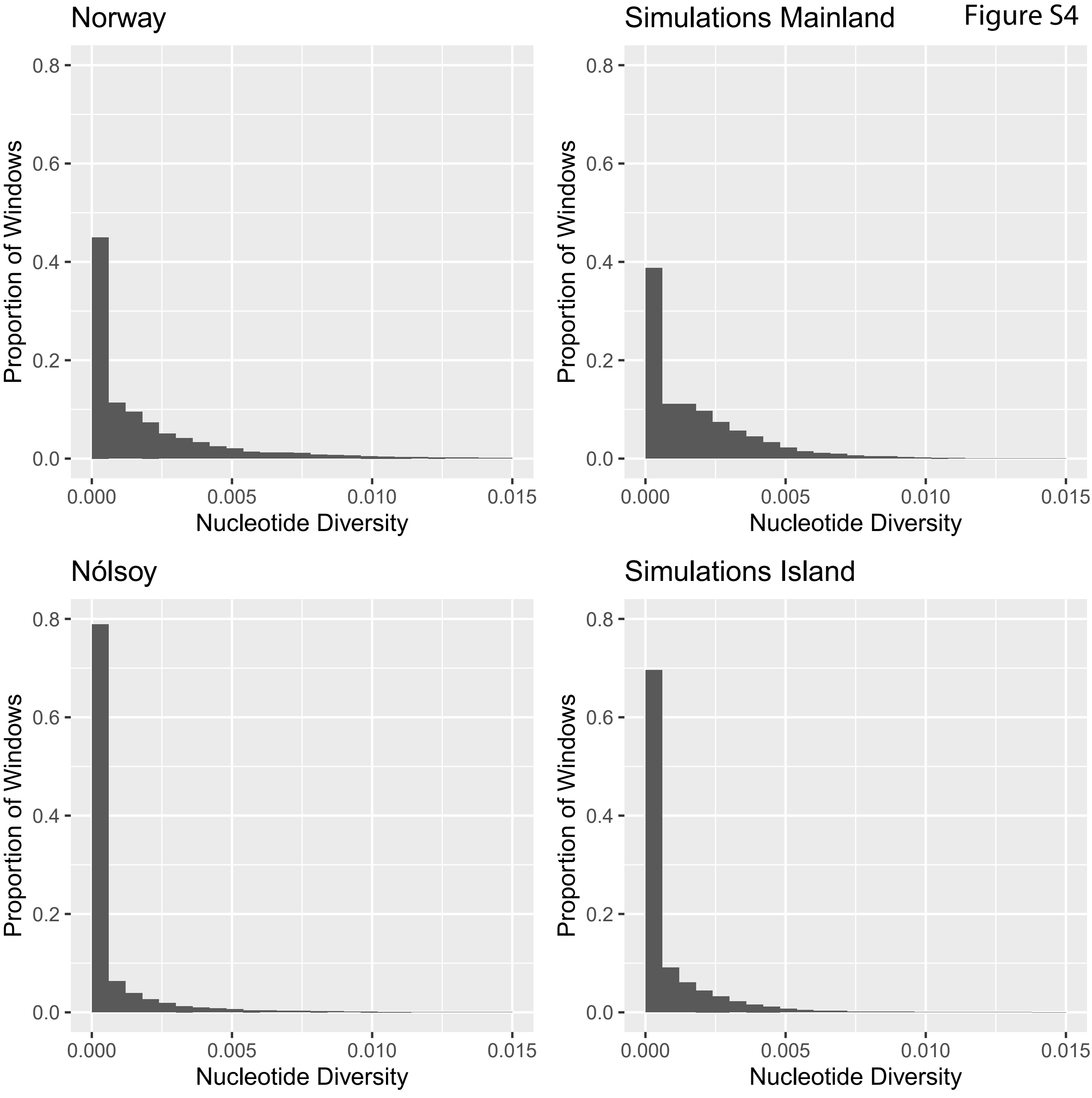

Norway
Simulations Mainland
Nólsoy
Simulations Island
Proportion of Windows
Nucleotide Diversity
0.0
0.2
0.4
0.6
0.8
0.000
0.005
0.010
0.015

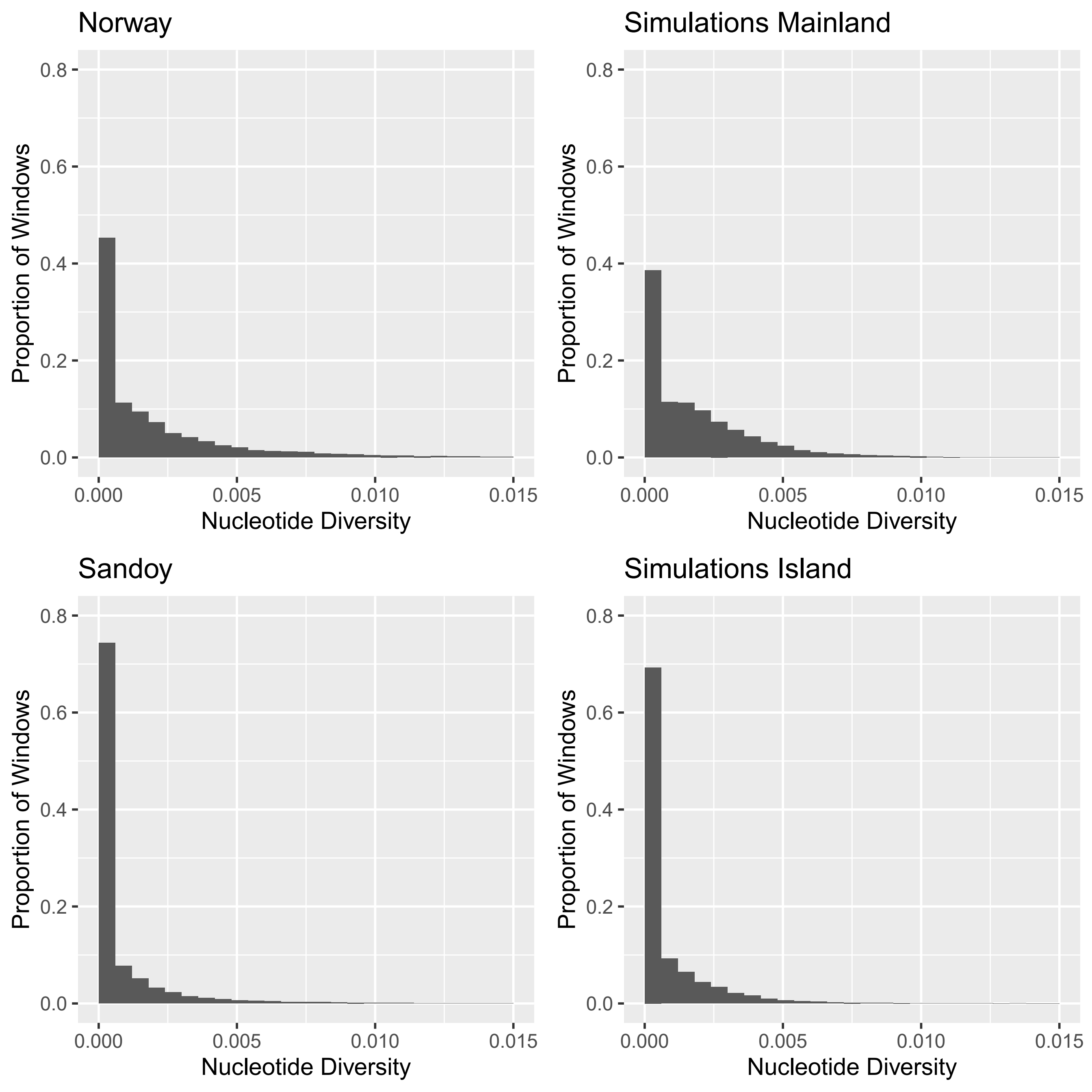

Norway
Simulations Mainland
Sandoy
Simulations Island
Proportion of Windows
Nucleotide Diversity
0.0
0.2
0.4
0.6
0.8
0.000
0.005
0.010
0.015

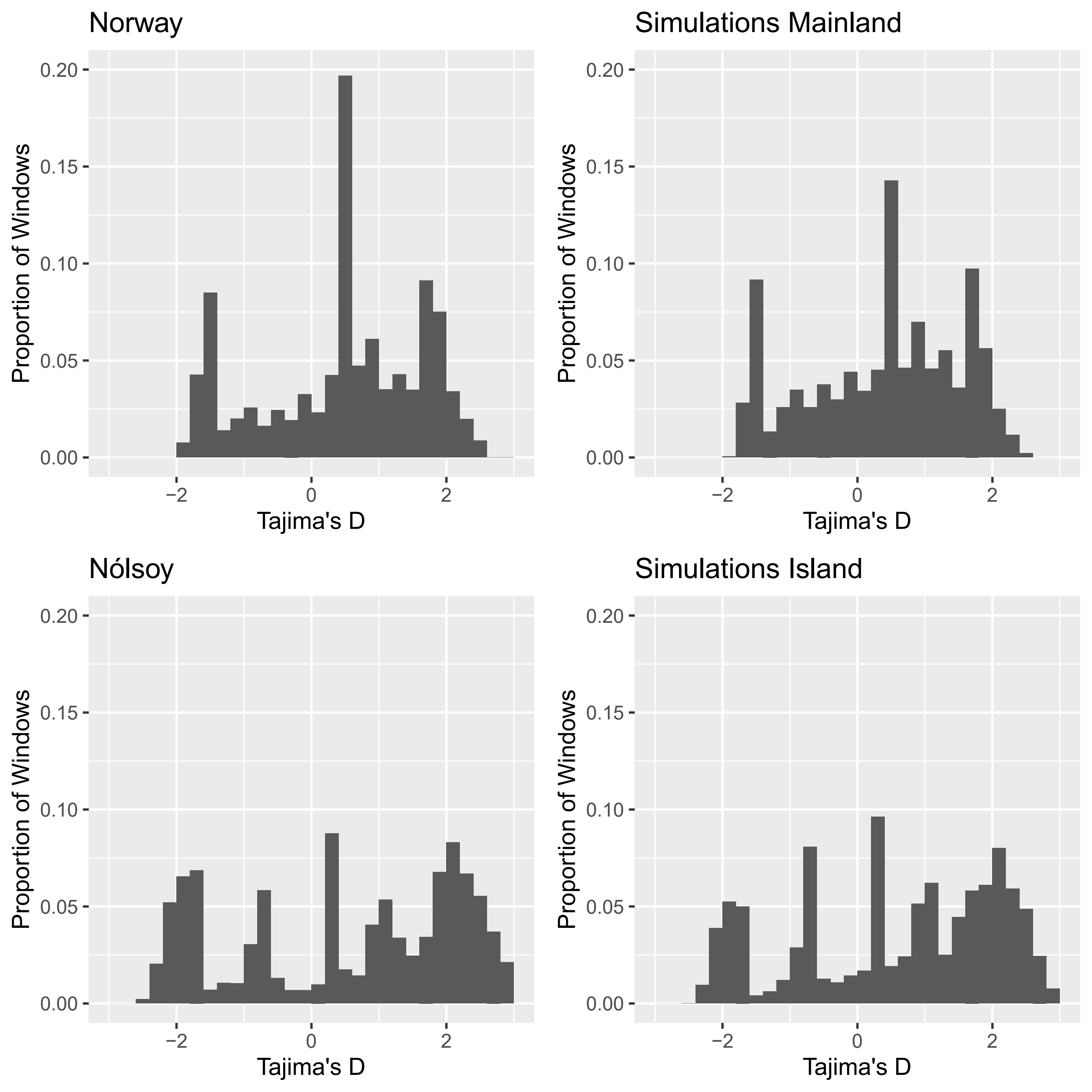

Norway
Simulations Mainland
Nólsoy
Simulations Island
Proportion of Windows
Tajima's D
0.00
0.05
0.10
0.15
0.20
−2
0
2

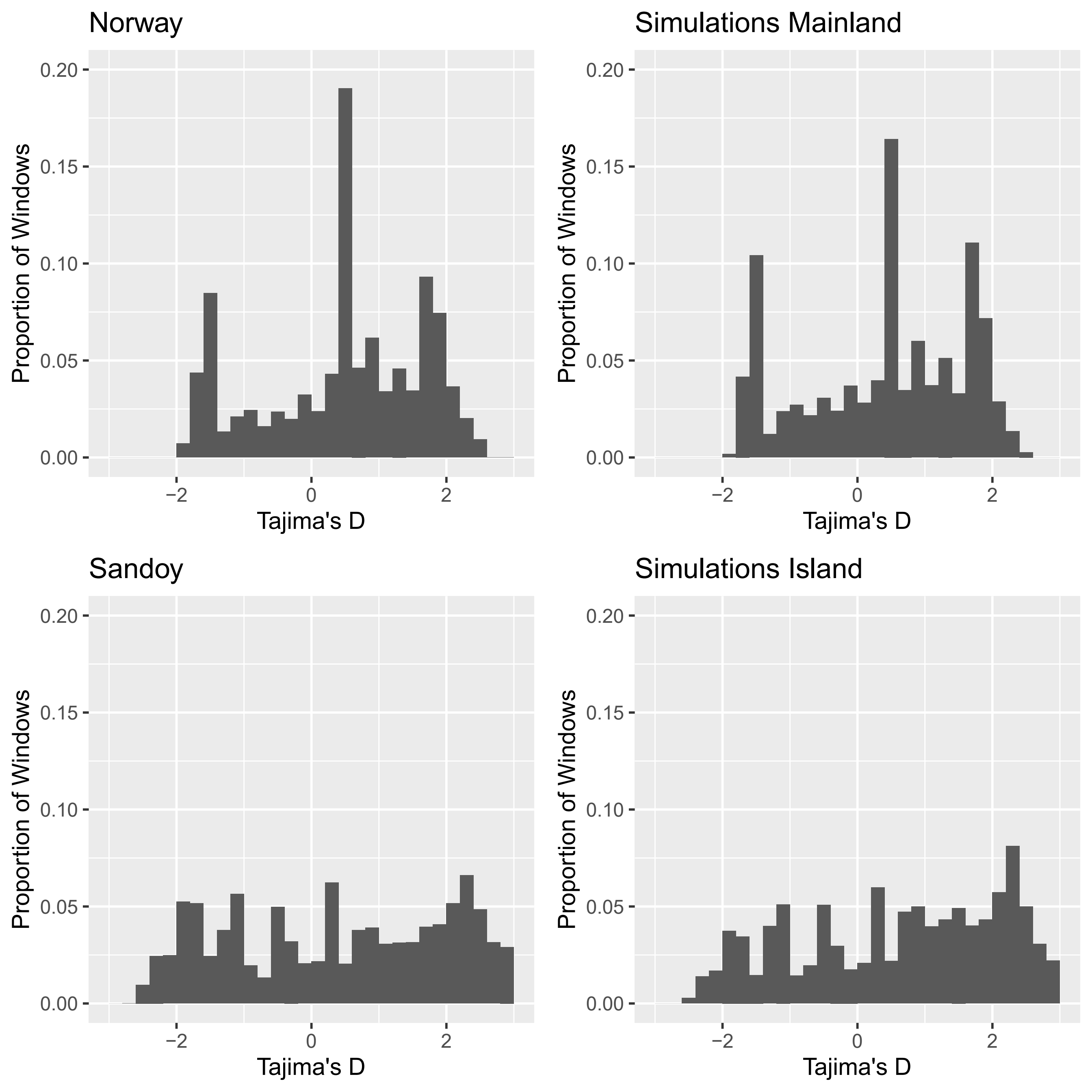

Norway
Simulations Mainland
Sandoy
Simulations Island
Proportion of Windows
0.20
0.15
0.10
0.05
0.00
−2
0
2
Tajima's D

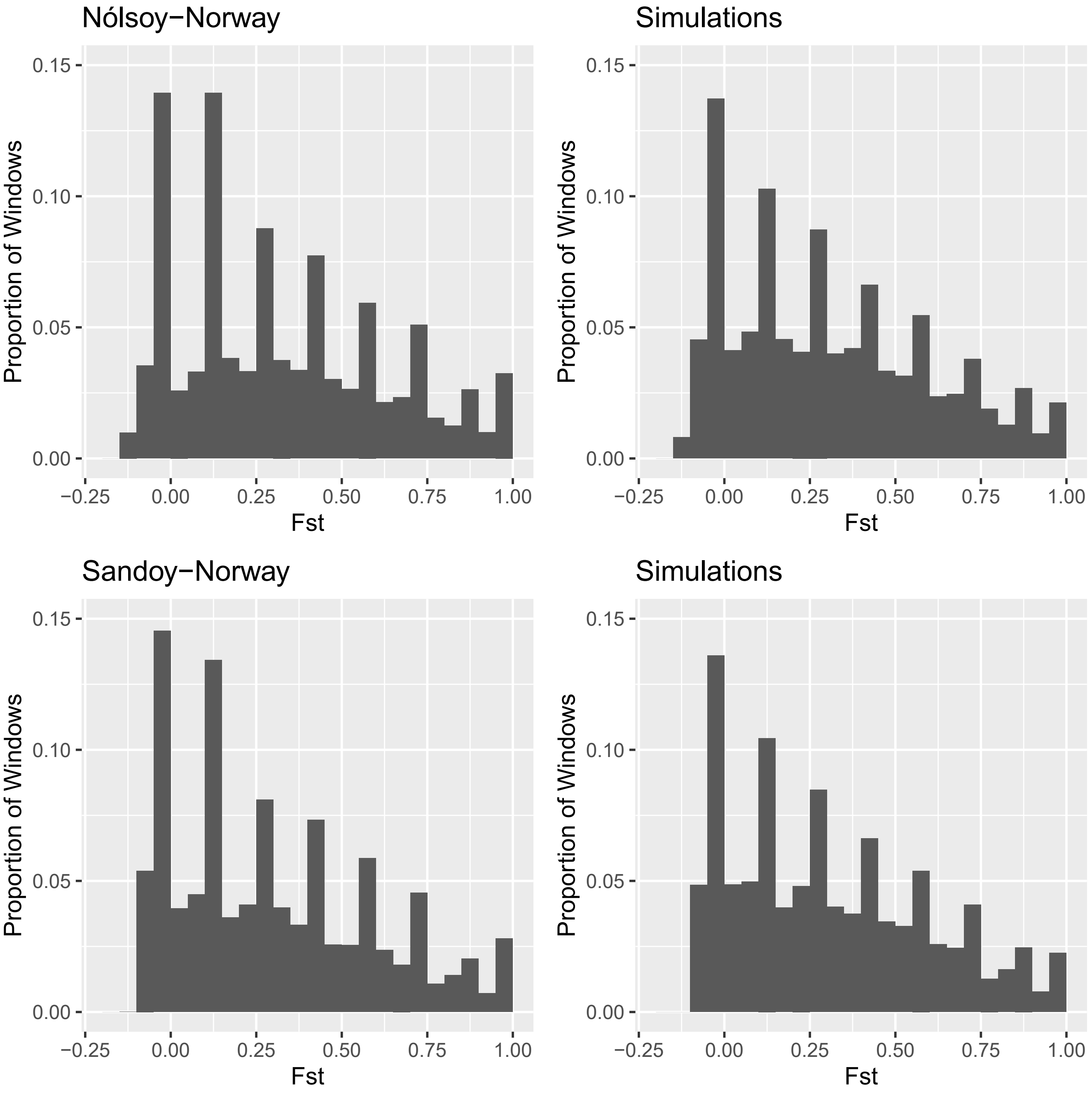
Nólsoy−Norway
Simulations
Sandoy−Norway
Simulations
Proportion of Windows
Fst
0.00
0.05
0.10
0.15
−0.25
0.00
0.25
0.50
0.75
1.00
